# Mangrove endophytes shorten the life cycle of rice while enhancing yield and salt tolerance

**DOI:** 10.1101/2024.07.12.603329

**Authors:** Amal Alghamdy, Anamika Rawat, Sabiha Parween, Arun Prasanna Nagarajan, Maged M. Saad, Heribert Hirt

## Abstract

Global climate change increasingly challenges agriculture with flooding and salinity. Among strategies to enhance crop resilience to these stresses, we tested several endophytic bacterial strains from mangroves, which are permanently exposed to flooding and high salinity. We show several strains that can enhance flooding and salinity tolerance in Arabidopsis and rice plants. Two strains and their combination massively enhanced the growth and yield of *Oryza sativa* cv. Nipponbare under both soil and hydroponic growth conditions with and without salt treatment. The bacteria-induced transcriptome changes in O. sativa roots related to ABA-signaling with lignin and suberin deposition in root tissues explain the altered responses of colonized rice plants to hypoxic and saline stress conditions. While enhancing yield and grain quality, bacterially colonized rice plants also show much earlier flowering, thereby massively shortening the life cycle of rice plants and opening the possibility for an additional harvest per year. These results show that microbes can be a powerful tool for enhancing the yield and resilience of rice to hypoxic and saline stress conditions.

## Introduction

Due to global warming, agriculture is challenged by the rising incidence of adverse weather events. Extreme events that alter water availability, like droughts and floods, threaten food security [1]. On a global scale, floods caused almost two-thirds of all damage and loss to crops between 2006 and 2016, costing billions of dollars [2].

Excessive water supply induces hypoxia in plants [3], increases the vulnerability to pathogen attack [4], and limits light flow to the plant [5]. During recovery after flooding, plants experience oxidative stress [6] and must remobilize nutrients to achieve a normal homeostatic state [7]. In recent years, flooding stress and its derivatives, like submergence, waterlogging, hypoxia, and anoxia, were investigated extensively in plants to identify molecular elements that may play a role in tolerance to flooding [8]. Plants respond to flooding and the associated stress by gene expression changes regulated at multiple levels, including transcription [3, 9] translation [10] and epigenetics [11].

Salinity is another global problem for plants that is becoming increasingly important. Soil salinity inhibits plant growth and development, strongly reducing crop yield. Salt stress induces both osmotic changes and toxicity in plants, resulting in reduced water availability and sodium accumulation [12]. Halophytes are plants adapted to saline soils using specialized strategies to deal with salinity. Since most crop species are salt-sensitive, efforts are underway to make them more salt-tolerant to maintain crop yield and potentially use saline irrigation.

The use of harmful chemical fertilizers and pesticides has negative impacts on the environment and human health. This situation has generated increasing interest in using beneficial microorganisms to develop sustainable agri-food systems [13]. Recent research showed the remarkable importance and range of microbial applications for enhancing the growth and health of plants, and plant growth-promoting bacteria (PGPBs) rapidly developed as a means to enhance tolerance to abiotic and biotic stresses [14]. So far, however, no microbes have been characterized that can enhance flooding tolerance and ensure the productivity of crops.

In this work, we used bacteria derived from the mangrove *Avicennia marina,* isolated from the shore of the Red Sea, whose roots are constantly submerged but also undergo repetitive flooding at the whole plant level. In addition, *A. marina* has adapted to grow under high saline conditions. We, therefore, tested several *A. marina* microbes first on *Arabidopsis thaliana* under waterlogging and then under waterlogging saline conditions. Selected single and multiple strains were then tested on *Oryza sativa cv.* Nipponbare under various conditions. Overall, our results identified strains that can protect rice plants from root submergence and saline stress. In addition, we could massively reduce the life cycle of rice, potentially allowing a shift from two to three harvests per year under optimal conditions.

## Methods

### Bacterial culture conditions

The bacteria used in this study, including *Isoptericola* sp. AK164 [15] and *Tritonibacter mobilis* AK171 were isolated from the rhizosphere of *Avicennia marina* seedlings growing on the Red Sea shore in 2019. They have been routinely grown in Zobell marine agar and incubated aerobically at 30 °C. The seeds were inoculated with either the two bacterial candidates individually (AK164 and AK171) or as a combination to create a synthetic community (SynCom). The inoculum of each bacterial strain used for successive experiments was 100 µL of O.D. 0.2 (∼10^9^ CFU mL^-1^ using a spectrophotometer at 600 nm). For the SynCom, a mixture of each strain was mixed as 50 μL of each (1:1).

### Plant material and growth conditions

The five-days-old *A. thaliana* seedlings, grown on ½ MS agar either non-inoculated or inoculated with single bacterial strains(s), were transferred on the top of a submerged ½ MS agar block (disk) to support the seedlings, in which a 24-well plate was filled with 500 μL of liquid media (Fig. S1). Control uninoculated seeds (MOCK) were grown in parallel.

For rice (*Oryza sativa* cv. Nipponbare), seeds were surface sterilized in 70% Ethanol with 0.05% Tween, rinsed with sterile distilled water five times, and then soaked in 50% sodium hypochlorite solution for 45 min. The seeds were rinsed well with sterile distilled water and transferred aseptically to Phytatray II (Sigma-Aldrich) containing ½ MS + 0.4% Agar. The trays were then kept in the dark at 30°C for 48 h to ensure synchronization in seed germination. The plants were grown in growth chambers at 27/25°C (day/night) with a 12:12 h light/dark cycle, the light intensity of 200 µmol photons m−2 s−1, and 70% humidity.

For RNASeq analysis, the 5-days-old seedlings, grown as described above, were transferred to buckets containing Hoagland solution supplemented with 0 or 100 mM NaCl. After 24h, the shoots and roots were collected separately and flash-frozen in liquid N_2_ before RNA extraction.

### Hydroponic growth conditions of *Oryza sativa* cv. Nipponbare

The potential plant growth-promoting (PGP) ability of 8 bacterial strains was tested hydroponically. The inoculum, as described above, was mixed with 50 mL of the ½ MS + 0.4% Agar. On day 5, the seedlings were transferred into 4.3 L buckets filled with half-strength modified Hoagland nutrient solution (5.6mM NH_4_NO_3_, 0.8mM MgSO_4_.7H_2_O, 0.8mM K_2_SO_4_, 0.18mM FeSO_4_.7H_2_O, 0.18mM Na_2_EDTA._2_H2O, 1.6mM CaCl_2_.2H_2_O, 0.8mM KNO_3_, 0.023mM H_3_BO_3_, 0.0045mM MnCl_2_.4H_2_O, 0.0003mM CuSO_4_.5H_2_O, 0.0015mM ZnCl_2_, 0.0001mM Na_2_MoO_4_.2H_2_O and 0.4mM K_2_HPO_4_.2H_2_O, pH of 5.8) (Fig. S2). To avoid contamination and to maintain continuous nutrient supply, the solution was changed every 3 days. For saline hydroponic growth, the concentration of NaCl was gradually increased until it reached 100 mM NaCl (1^st^ week 0 mM → 2^nd^ week 50 mM → 3^rd^ week 75 mM → 4^th^ week 100 mM). A total of 34 plants were used for each treatment (Fig. S2). After 28 days, the plants were scored for their phenotypes.

### Greenhouse experiments

The experiments were conducted in the greenhouse facility at KAUST from February 2022 till September 2023. The potting soil (Stender) mixed with rocks (3:1) was used in this study. The 5-days-old rice seedings mock-inoculated or inoculated with either AK164, AK171, or the SynCom (AK164 +AK171, 1:1) were transferred to pots and irrigated with Hoagland solution (0 mM NaCl) or Hoagland solution supplemented with NaCl, gradually increasing the NaCl concentration to 100 mM for the duration of one month (Fig. S2).

After the first month, all plants were irrigated with tap water till flowering and stopped when tillers were filled with seeds. The panicles were then ripened on the plant until harvest. After 90 days, the plant phenotypic data were collected, including plant height, number of panicles, and spikelet per plant. Two weeks later, the plants were harvested from the pots, and selected parameters were measured, including plant height (cm) and root and shoot DW (gm). Other phenotypic parameters included the number of tillers, the number and weight of panicles per plant, the grain yield as well as the number and weight of grains per plant.

### RNA-Seq Library Preparation and Transcriptome Analysis

Total RNA was extracted from roots and shoots of 21-day-old mock and AK164, AK171 colonized plants using Direct-zol RNA Miniprep Plus Kit following the manufacturer’s instructions (ZYMO RESEARCH; USA). The quality and quantity of the RNA were assessed by Agilent 2100 Bioanalyzer (RNA integrity number >7) and Qubit TM 3.0 Fluorometer with TNA BR assay kit. Samples were processed further for library preparation, sequencing, and bioinformatic analysis (Novogene Co., LTD (Beijing, China). The RNASeq data have been submitted to the public data base GEO and were assigned GSE269165. Genes with adjusted p-value < 0.01 and |log2 (FoldChange)| > 1 were considered differentially expressed compared to control. The following secure token has been created to allow review of the transcriptome data: **mhgjmcqcvfkhpch.**

The reference genome *O. sativa* Japonica cv. Nipponbare (IRGSP-1.0) and gene model annotation files were downloaded from RAP-DB [16, 17] (https://rapdb.dna.affrc.go.jp/). Hisat2 v2.05 was used to align the RNAseq reads with the reference genome to generate a database of splice junctions on the gene models. Next, StringTie (v.3.3b) [18] was used to assemble and quantify full-length transcripts representing multiple splice variants for each gene locus and feature Counts v1.5.0-p3 to count the read numbers mapped to each gene. Then, the FPKM of each gene was calculated based on the gene length and read count mapped to this gene. We then used <u>R</u>. Further, the enrichment analyses were performed using cluster Profiler R package with corrected p-value cut-off < 0.05 to call out statistical significance. Gene Ontology (GO) enrichment analysis of differentially expressed genes was implemented by the DESeq2, in which gene length bias was corrected. GO terms with corrected *p-value* less than 0.05 were considered significantly enriched by differential expressed genes. KEGG was used for functions and utilities of the biological system (http://www.genome.jp/kegg/).

### Histochemical staining of rice roots

Roots of 16-day-old rice plantlets were fixed in fixative (Formaldehyde: Ethanol: Acetic Acid, 10%:50%:5%), and the semi-thin sections were cut from the zone 3 cm above the root tip. Briefly, the root samples were embedded in 5% low melting agarose, and 100-um thick sections were cut with the help of a vibratome (Leica VT1200S). The sections were stained with Calcofluor white, Fluorol Yellow, and Basic Fuchsin before being visualized using Zeiss LSM880 confocal microscope as described in Sexauer et. al. [19].

### Statistical Analysis

The data from the plant screening assay were subjected to non-parametric one-way ANOVA and the Kruskal-Wallis test [20]. The statistical difference is based on Dunn’s multiple comparison tests. All statistical analysis was done using GraphPad Prism version 9.5.0 (525) software (https://graphpad.com).

## Results

### Mangrove bacteria enhance the resilience of *Arabidopsis thaliana* to salinity under waterlogging conditions

To mimic the hypoxic conditions naturally occurring in the mangrove ecosystem, *A. thaliana* seedlings were inoculated with bacterial strains and grown by a waterlogging method (Figure S1). Based on previous analyses (unpublished data), 16 isolates were tested for their ability to enhance plant growth on ½ MS in waterlogging growth conditions. When compared to non-inoculated controls, five strains (AK031, AK116, AK164, AK171, AK225) showed significant enhancement (35 ‒107 %) of plant growth under both waterlogging (1/2MS) and saline waterlogging conditions (1/2MS + 100 mM NaCl) (Table 1, Figure 1).

**Figure 1.**
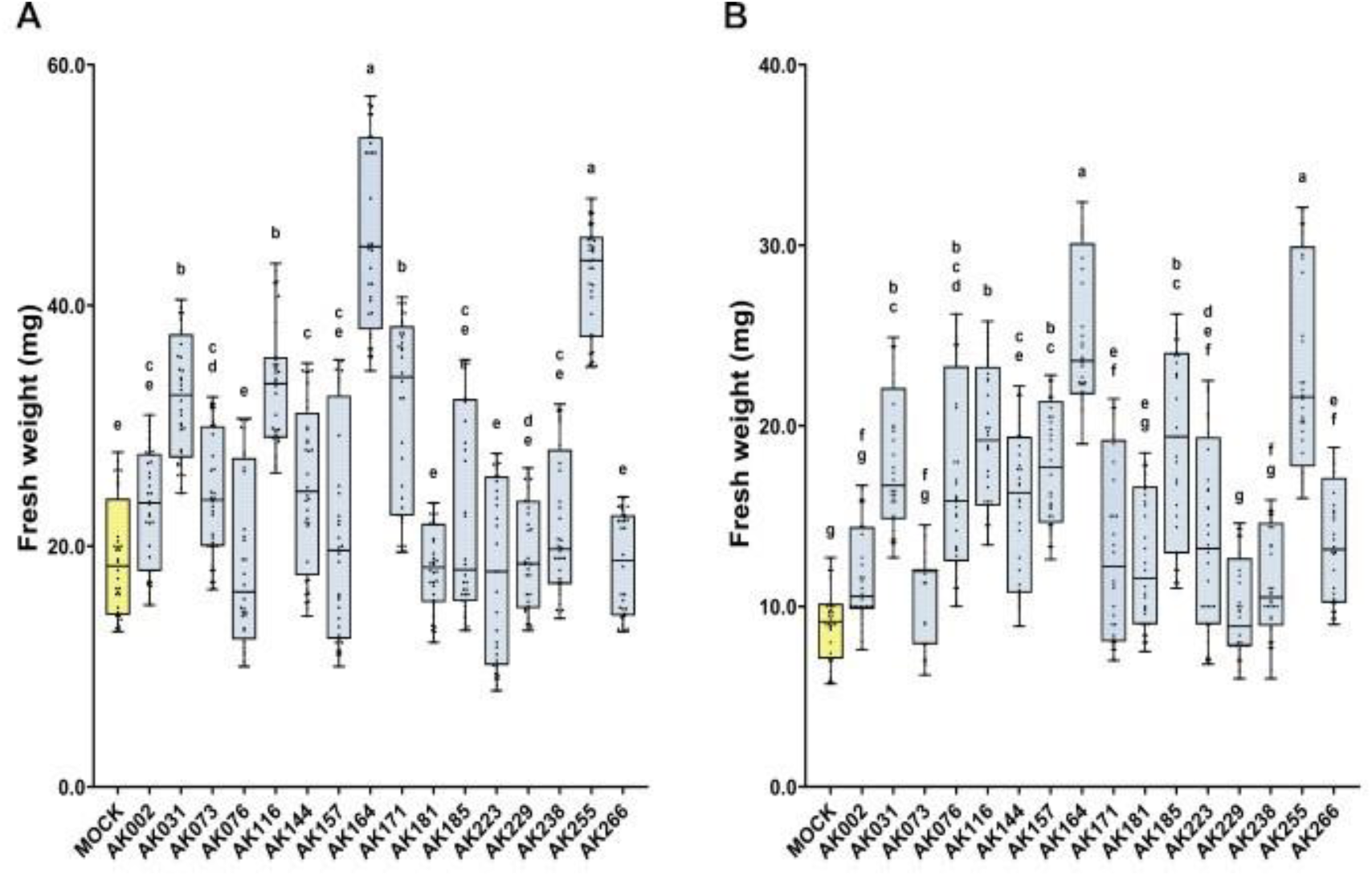
Mangrove strains enhance the growth of A. thaliana under waterlogging conditions. Growth enhancement of *A. thaliana* seedlings inoculated with candidate bacteria compared to non-inoculated seedlings (MOCK) in the waterlogging method. Box plots represent FW of *A. thaliana* grown on A-½ MS, B-½MS + 100mM NaCl. The statistically significant differences based on ANOVA followed by post hoc Tukey’s analysis are represented as a compact letter display of all comparisons (P < 0.05).

**Table 1:** Beneficial increase (%) of *A. thaliana* fresh weight (gm) by 16 candidate strains under submergent growth conditions. p>0.05, **P>0.01, *** P>0.001, **** P>0.0001.

| No. | Strain | Name | ½ MS | ½ MS<br>+ 100mM<br>NaCl |
| --- | --- | --- | --- | --- |
| 1 | AK002 | *Paenibacillus lautus* (OR447938) | 9.5 | 21.8 |
| 2 | <u>AK031</u> | *Bacillus seohaeanensis* (OR447774) | 45.8**** | 96.1*** |
| 3 | AK073 | *Thalassospira tepidiphila* (OR447979) | 12.8 | 24.4 |
| 4 | AK076 | *Tritonibacter mobilis* (OR447785) | -8.2 | 87.3**** |
| 5 | <u>AK116</u> | *Halobacillus locisalis* (OR447813) | 43.0*** | 125.2*** |
| 6 | AK144 | *Microbulbifer elongatus* (OR447909) | 17.5* | 59.7** |
| 7 | AK157 | *Demequina activiva* (OR447779) | -1.9 | 78.5**** |
| 8 | <u>AK164</u> | *Isoptericola chiayiensis* (OR447867) | 107.0**** | 165.3**** |
| 9 | <u>AK171</u> | *T. mobilis* (OR447787) | 35.2* | 47.3* |
| 10 | AK181 | *Pseudomonas azotoformans* (OR447934) | -20.7 | 57.4* |
| 11 | AK185 | *T. mobilis* (OR447793) | 1.6 | 102.5**** |
| 12 | AK223 | *P. azotoformans* (OR447928) | -10.1 | 58.0* |
| 13 | AK229 | *Martelella mangrovi* (OR447916) | -3.1 | 11.2 |
| 14 | AK238 | *Celerinatantimonas diazotrophica* (OR447776) | 6.9 | 17.9 |
| 15 | <u>AK255</u> | *I. chiayiensis* (OR447854) | 101.4**** | 146.1**** |
| 16 | AK266 | *P. lautus* (OR448017) | 9.5 | 48.7** |

### Mangrove bacteria enhance the resilience of *O. sativa* cv. Nipponbare using hydroponics

To test for the beneficial effects of the *A. marina bacterial* strains on rice, we tested eight selected strains on rice grown in hydroponic as well as hydroponic saline conditions. Among the eight tested strains (AK031, AK073, AK116, AK157, AK164 [15], AK171, AK181, AK255), several exhibited notable growth enhancements in inoculated rice plants compared to the non-inoculated plants under the hydroponic conditions (Figure 2 A-C), Figure S3). Under saline hydroponic conditions (100 mM NaCl), two strains (AK164 and AK171) not only promoted growth (Figures 2 D and E; Figure S4) but also significantly increased the number of tillers per plant in inoculated rice relative to non-inoculated plants (Figure 2 F).

**Figure 2:**
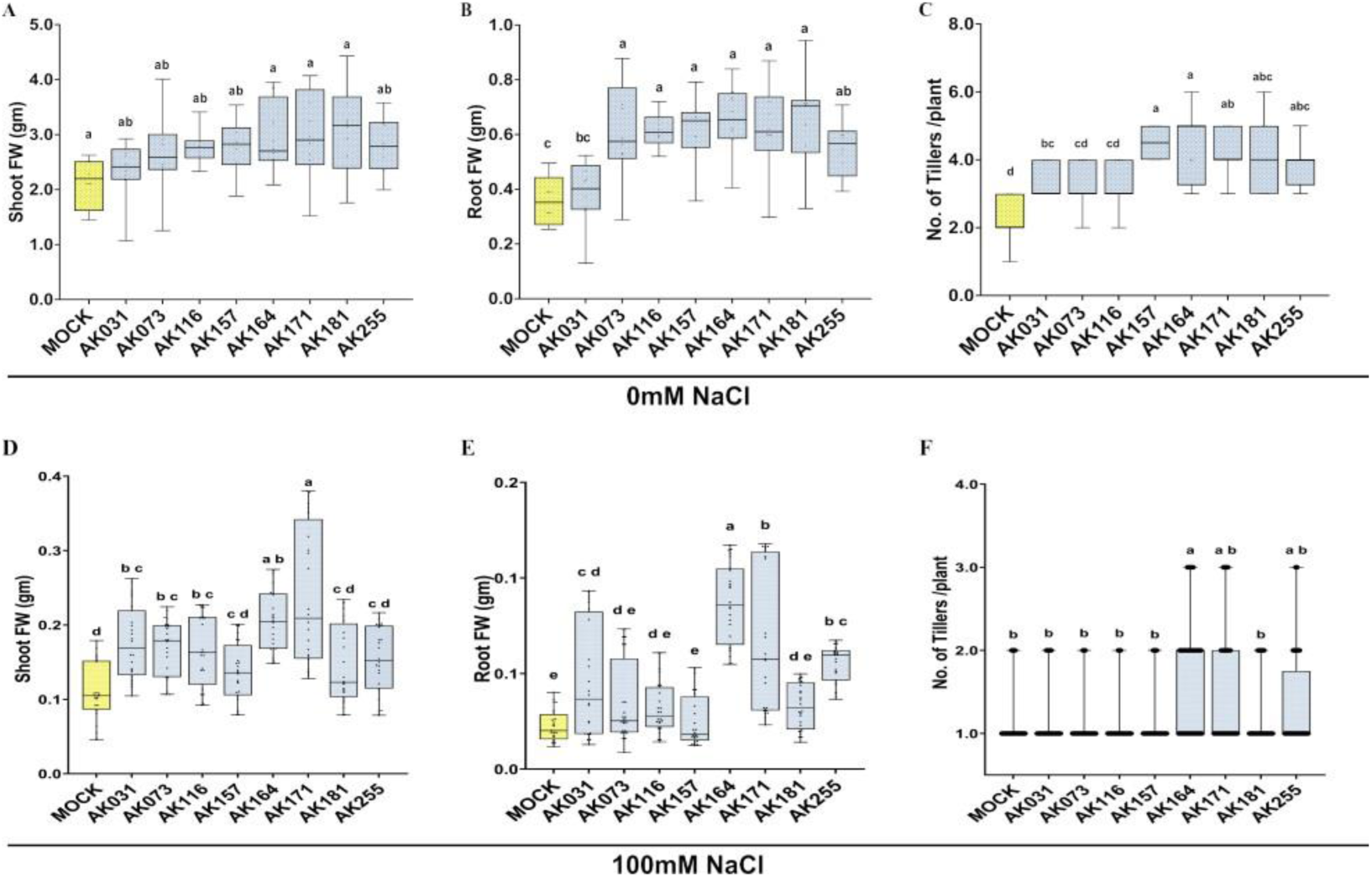
Mangrove bacteria enhance the growth of rice. Hydroponically cultivated rice *O. sativa* cv. Nipponbare was analyzed at 28 days post-inoculation with different bacterial strains or mock-inoculation at either 0 mM NaCl (A-C) or 100 mM NaCl (D-F). A, D-shoot fresh weight, B, E-root fresh weight, C, F-number of tillers per plant. The statistically significant differences based on ANOVA followed by post hoc Tukey’s analysis are represented as a compact letter display of all comparisons (P < 0.05).

### AK164, AK171, and SynCom enhance rice growth under hydroponic normal and salt stress conditions

AK164 and AK171 were next analyzed alone and in combination (SynCom) for their potential to enhance the growth and salt stress resistance of rice. As described beforehand, we used a hydroponic system to assess various growth enhancement parameters in hydroponic normal conditions (Figure 3 A-D). To evaluate the efficacy, we assessed various growth parameters, including dry weight (DW) and the number of tillers, compared to non-inoculated plants (MOCK).

**Figure 3.**
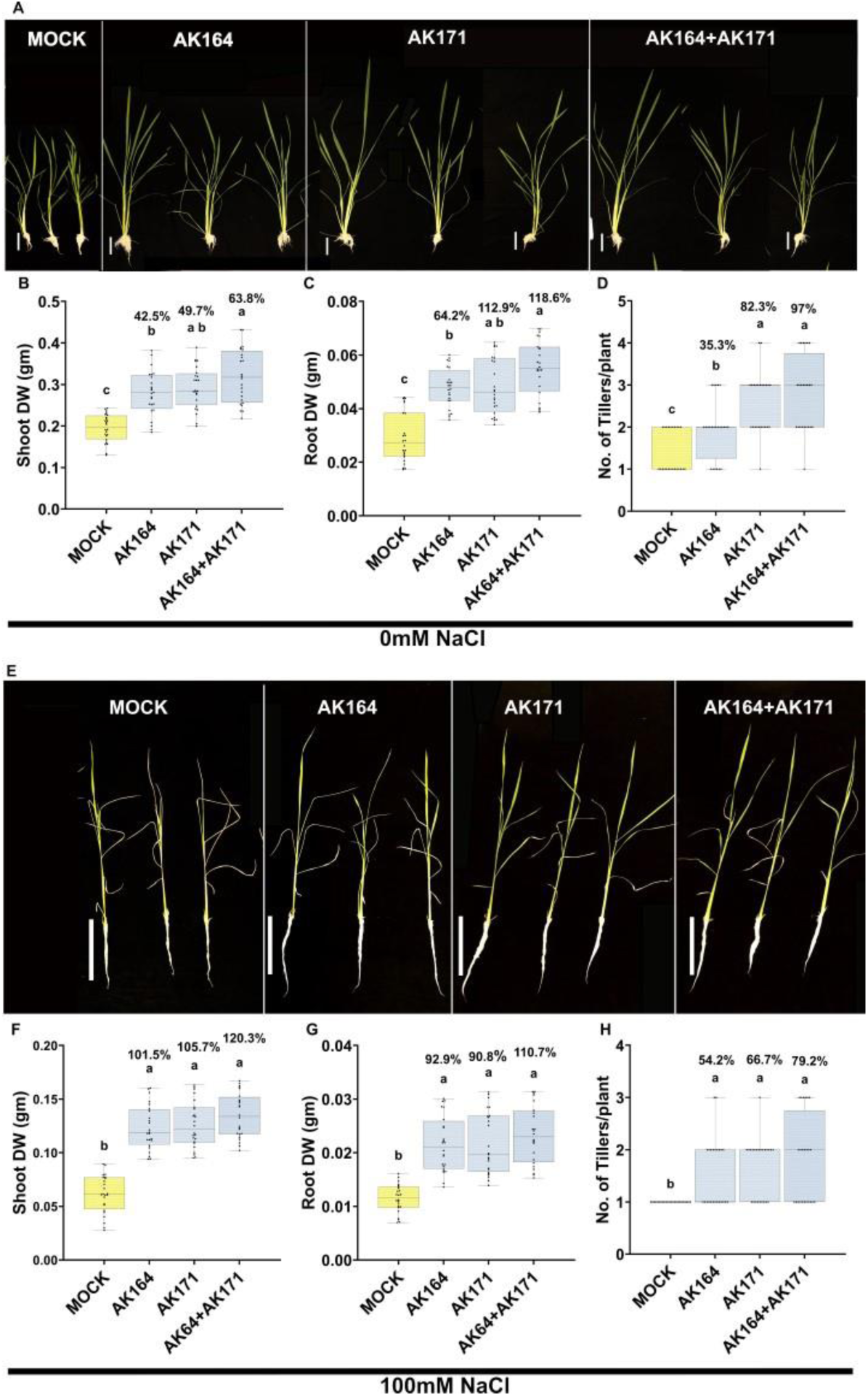
AK164, AK171, and SynCom enhance rice growth under hydroponic normal and hydroponic saline conditions. A, E-Plant phenotypes of hydroponically or hydroponically saline-grown rice at 28 days post-inoculation with MOCK, AK164, AK171, or SynCom. Bar = 10 cm. Percent beneficial increase in growth and number of tillers under hydroponic normal or hydroponic saline conditions of inoculated compared to non-inoculated (MOCK) plants: B, F-shoot dry weight (gm), C, G-root dry weight (gm), D, H-number of tillers per plant. The statistically significant differences based on ANOVA followed by post hoc Tukey’s analysis are represented as a compact letter display of all comparisons (P < 0.05).

Both AK164 and AK171 significantly increased the shoot DW of inoculated rice plants by 42.5% and 49.7%, the root DW by 64.2% and 112.6%, and the number of tillers by 35.3% and 82.3%, respectively, when compared to mock-inoculated plants (Figures 3 A‒D). Subsequently, we investigated the combined effects of these two strains as a SynCom, which resulted in a considerably higher DW in both shoots and roots when compared to the single inoculated strains, resulting in 63.8% and 118.6% increase, respectively, compared to mock-inoculated plants (Figures 3 B and C, Figure S5). Importantly, the tiller number per plant for SynCom inoculated rice increased from 35.3% (AK164) and 82.3% (AK171) to 97% (AK164+AK171), as shown in Figure 4 D.

**Figure 4.**
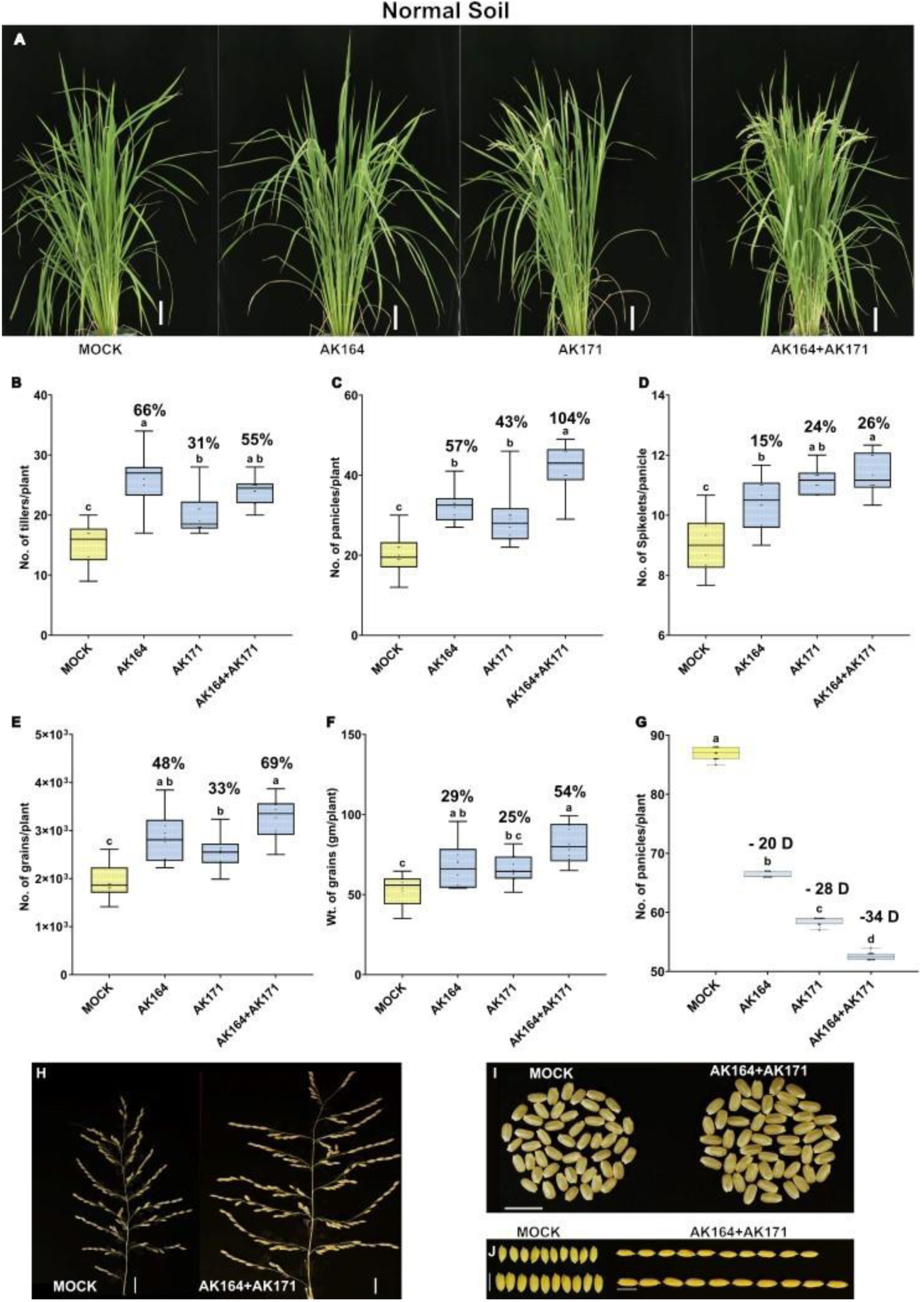
AK164, AK171, and SynCom enhance the growth and yield of normal soil-grown rice. Growth of *O. sativa* cv. Nipponbare inoculated with AK164, AK171, SynCom (AK164+AK171), or MOCK on normal soil (0 mM NaCl). A-Rice phenotypes at 90 days of growth (Bar = 10 cm). B to G Percent beneficial increase in the number of tillers per plant (B), number of panicles per plant (C), number of spikelets per panicle (D), number of grains per plant (E), weight of grains per plant (F), and flowering onset (G) of AK164, AK171 or SynCom inoculated plants compared to non-inoculated (MOCK). The statistically significant differences based on ANOVA followed by post hoc Tukey’s analysis are represented as a compact letter display of all comparisons (P < 0.05). H-Comparison of panicle size of mock (top row) and SynCom (bottom row) inoculated rice (Bar = 2 cm). I-50 seed size (Bar = 10 mm) and J-10 seed length (Bar = 7 mm).

Under hydroponic saline conditions (100mM NaCl) (Figure 3 E-H), rice plants inoculated with AK164, AK171, or SynCom exhibited significant enhancements in growth compared to the mock (Figure 3 E). Notably, the SynCom inoculation showed substantial enhancement in dry weight in both shoot (up to 120%) and root (up to 110%). Also, the number of tillers per plant was higher (up to 79%) than the mock. These findings underscore the potential of AK164 and AK171, both individually and in combination, to promote plant growth and mitigate the effects of abiotic stress.

### AK164 and AK171 enhance soil-grown rice performance and yield

We then conducted greenhouse experiments (Figure S2A) to evaluate the growth of rice seeds inoculated with individual bacterial strains (AK164 and AK171) and their SynCom with/without salinity treatments. After 90 days of growth, we analyzed various agronomic parameters to assess the effects of bacterial inoculation on rice growth (Figures 4 and 5, Figures S6 and S7). Under non**-** stressed conditions (0 mM NaCl) as shown in Figure 4 A), we observed that AK164 or AK171 alone already substantially improved most agronomic parameters (Figures 4 B-J). Significant increases were observed in the production of the number of tillers/plant (31-66%), panicles/plant (43-104%), and spikelet/panicle (15-16%). As before, SynCom-treated rice showed superior PGP metrics in all these categories. Most importantly for yield assessment, the number and weight of grains per plant also showed increases of 48% and 29% for AK164, 33% and 25% for AK171, and 69% and 54% for SynCom treated rice plants, respectively (Figure 4 I – J).

**Figure 5.**
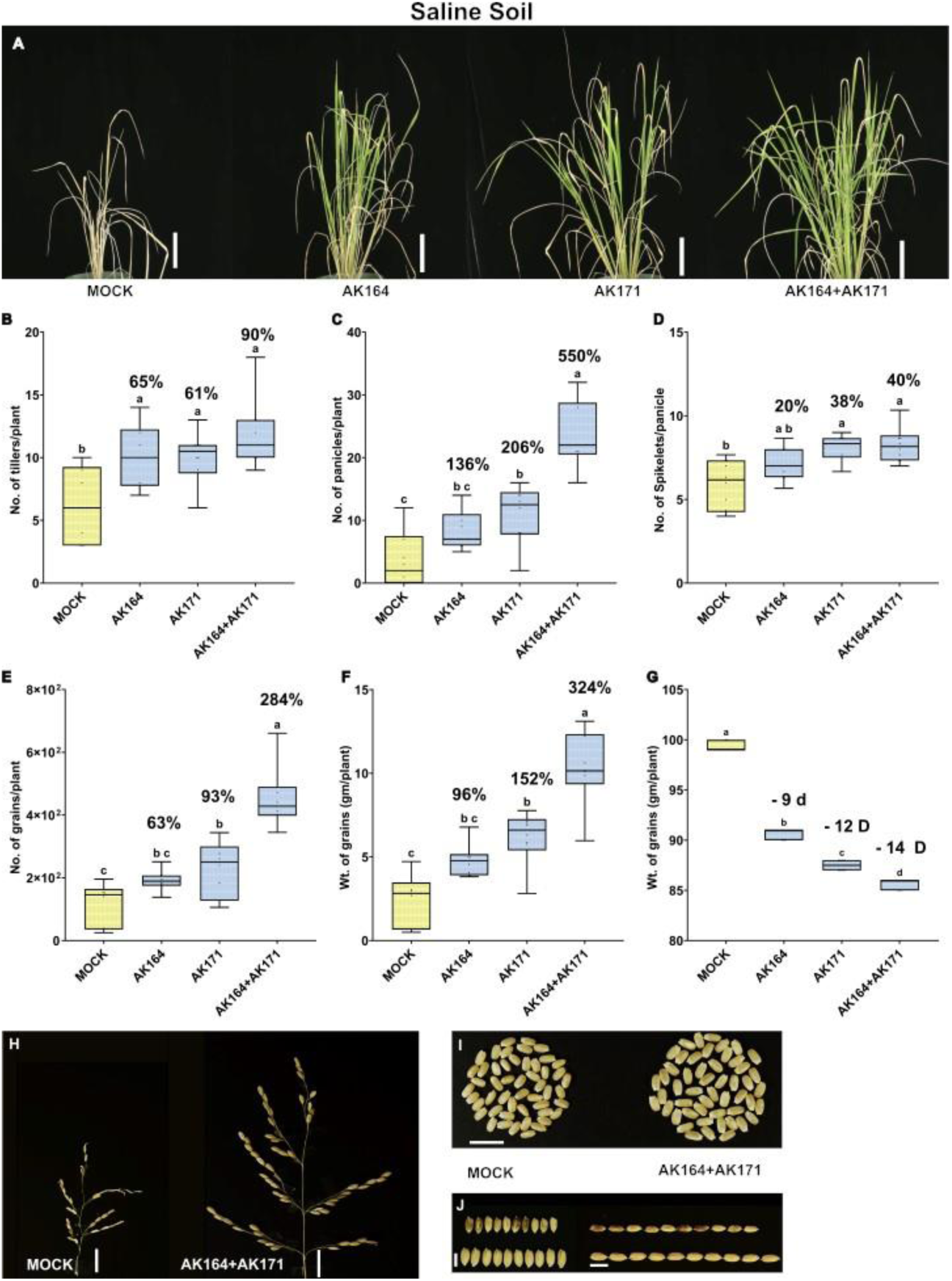
AK164, AK171, and SynCom enhance growth and yield of saline soil-grown rice. Growth of *O. sativa* cv. Nipponbare inoculated with AK164, AK171, SynCom (AK164+AK171), or MOCK on saline soil (100 mM NaCl). A-Rice phenotypes at 90 days of growth (Bar = 10 cm). B to G Percent beneficial increase in the number of tillers per plant (B), number of panicles per plant (C), number of spikelets per panicle (D), number of grains per plant (E), weight of grains per plant (F), and flowering onset (G) of AK164, AK171 or SynCom inoculated plants compared to non-inoculated (MOCK). The statistically significant differences based on ANOVA followed by post hoc Tukey’s analysis are represented as a compact letter display of all comparisons (P < 0.05). H-J-Yield increase of SynCom inoculated rice on saline soil. H-Comparison of panicle size of mock (top row) and SynCom (bottom row) inoculated rice (Bar = 2 cm). I-50 seed size (Bar = 10 mm) and J-10 seed length (Bar = 7 mm).

Of particular interest was the observation that AK164 and AK171 accelerated flowering onset in rice plants. While non-inoculated rice plants flowered around 88 days, AK164 and AK171 advanced flowering by 20 and 28 days, respectively, after transfer to soil (Figure 4G). Remarkably, SynCom accelerated flowering by 34 days, shortening the entire life cycle of *O. sativa* from 4 to 3 months while simultaneously increasing the weight and number of grains per plant by 54% and 69%, respectively (Figure 4 G). Furthermore, both panicles and spikelet were clearly longer, which explains the significant enhancement of the weight of panicles per plant in the plants inoculated with the SynCom compared to non-inoculated (mock) plants (27% and 31%), as shown in Figure 4 H. In addition, the size and length of seeds (Figures 4 I and J) were significantly enhanced, indicating the potential of bacterial inoculation to improve seed quality and yield in rice cultivation.

Under saline conditions (100 mM NaCl), the plants treated with either of the bacteria AK164 or AK171 alone did enhance the agronomic parameters (Figure 5 A). Still, SynCom notably showed a further increase in the number of tillers by 90% (Figure 5 B), panicles per plant by 550% (Figure 5 C), spikelet per panicle by 40% (Figure 5 D), as well as the number and weight of grains per plant (Figures 5 E and F). The accelerated flowering onset of rice was also observed for plants under saline conditions. While non-inoculated rice plants flowered around 100 days, AK164 and AK171 advanced flowering by 9 and 12 days, respectively (Figures 5 G). However, SynCom (AK164+AK171) accelerated the flowering of rice by 14 days.

The yield of AK164, AK171, or SynCom inoculated rice was remarkably enhanced as the weight and number of grains per plant were significantly increased by 324 and 284%, respectively (Figures 5 H-J). Both panicles and spikelet were clearly longer and more filled with seeds, which explains the significant enhancement of the weight of panicles per plant in rice inoculated with the bacterial strains when compared to non-inoculated plants (1032% and 4915%), as shown in Figure 5 H. In addition, the size and length of seeds (Figure 5 I and J) were significantly enhanced, indicating the potential of the bacteria to improve seed quality and yield in rice cultivation under saline conditions.

### AK164 and AK171 induce upregulation of rice ABA and carotenoid signaling pathways

To gain a better understanding of the molecular mechanisms by which these bacteria enhance the growth of the rice plants, we performed RNA seq analysis of AK164 and AK171 colonized rice plants with and without salt stress conditions. Since the mangrove bacterial strains are both root epiphytes, we conducted this analysis separately for rice root and shoot tissues and concentrated on the rice root transcriptome (Figures 6 A-C). For roots of AK164-colonized rice plants, 71 genes were found to be up-regulated (Figure 6 B), among which were the genes for the two carotenoid biosynthesis enzymes CCD1 and CCD8a, the SAPK9 protein kinase, the NRT1/AIT1 transporter, and the histone demethylase JMJ30 (Figure 6 C). For the 12 down-regulated genes (Figure 6 B), the only GO term was found for transporters, but almost exclusively contained aquaporin genes PIP1;2, PIP1;3, PIP2;A13, PIP2;3, PIP2;4, PIP2;5 and TIP2;1 (Figure 6 C). In roots of AK171-colonized plants, 107 genes were found to be up-regulated (Figure 6 B), comprising again JMJ30, SAPK9, PCL1, and NRT1, but also CCD1 and LOX1, LOX3 (Figure 6 C). Interestingly, in the 37 down-regulated genes, the only GO term with high significance (5×10E-5) was retrieved for transporters, most of which comprised the same set of aquaporin genes as found in the AK164-inoculated plants’ transcriptome, except for PIP1;2 (Figure 6 C). Although roots of non-colonized plants showed the regulation of massive sets of DEGs under saline conditions, AK164-or AK171-colonized plant roots showed no significant up or down-regulated genes (Figure 6 B).

**Figure 6:**
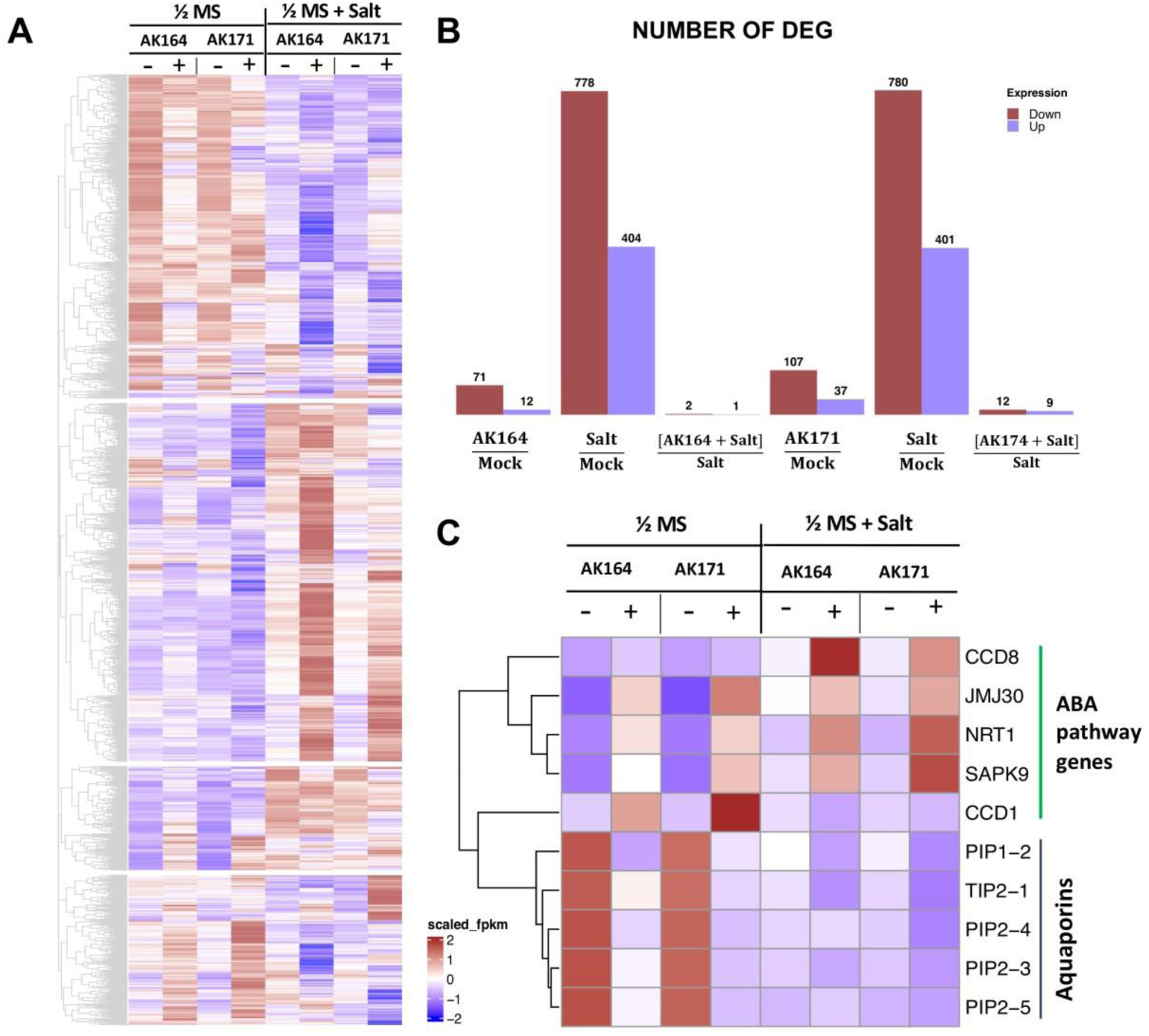
Heatmap of RNA sequencing data. DEGs of rice plants grown under normal and saline hydroponic conditions stress treated with AK164, AK171, or MOCK inoculated. A-Overview of DEGs in rice roots colonized with AK164, AK171, or MOCK under normal (0 mM NaCl) and saline (100 mM NaCl) hydroponic conditions. B-Number of DEGs in different conditions. C-Group of selected root DEGs (log 2 > 1, P < 0.01).

For shoots of AK164-colonized rice plants grown under normal hydroponic conditions, no GO terms were retrieved, neither for up or down-regulated genes (Figure S3). However, the upregulated shoot gene set in both AK171-and AK164-colonized plants contained the genes for JMJ30, SAPK9, NRT1, the transcription factor PCL1, and the receptor kinase Feronia in control conditions. For the down-regulated 171 genes, no GO terms were obtained. Under saline conditions, the shoot transcriptome of AK164-colonized plants only showed 12 up-and 69 down-regulated DEGs with no significant GO terms. However, SAPK9, NRT1, PCL1, and JMJ30 were again found in the significantly upregulated list of genes.

The high overlap between the root transcriptomes of AK164-and AK171-colonized rice plants was surprising and indicated that the two taxonomically different strains were targeting similar set of genes. Interestingly, a closer look at the genes revealed that most of these differentially expressed genes were either biosynthesis, signaling, or target genes of the carotenoid pathway (Figure 6 C). Intriguingly, some of the target genes, such as JMJ30, NRT1, and SAPK9, have been shown to be ABA-regulated. In contrast, the carotenoid biosynthesis genes CCD1 and CCD8 encode oxygenases, producing apocarotenoids including β-cyclocitral and strigolactone.

### AK164 and AK171 promote the deposition of secondary cell wall components in colonized rice

Aiming to visualize the secondary cell wall components in rice roots, we performed triple staining of rice roots grown hydroponically under normal conditions. Basic Fuchsin has been shown to stain lignin. In contrast, Fluorol Yellow stains suberin. The two dyes were combined to visualize the deposition of these two important components of the cell wall, which play important roles as a barrier in preventing loss of water and solutes and act to enhance plant cell wall rigidity. Rice tolerates hypoxia in the surrounding solution by forming lysigenous aerenchyma and a physical barrier to prevent radial oxygen loss (Figure 7). We observed changes in sclerenchyma layer formation in colonized plants. The sclerenchyma layer is a single-cell file in WT rice roots. While colonized plants showed multiple sclerenchyma layers, AK171 induced multi-serrate sclerenchyma layer formation, which was also seen in the AK164+AK171 inoculated plants. Multi-serrate sclerenchyma is associated with greater root lignin concentration and greater tensile strength for plants to survive in low-oxygen conditions. Deposition of suberin in endodermal cells is another adaptation in rice roots. We observed enhanced suberized endodermal cells in rice roots when colonized with PGPB candidates, suggesting enhanced barrier formation to prevent diffusion through plasma membranes.

**Figure 7.**
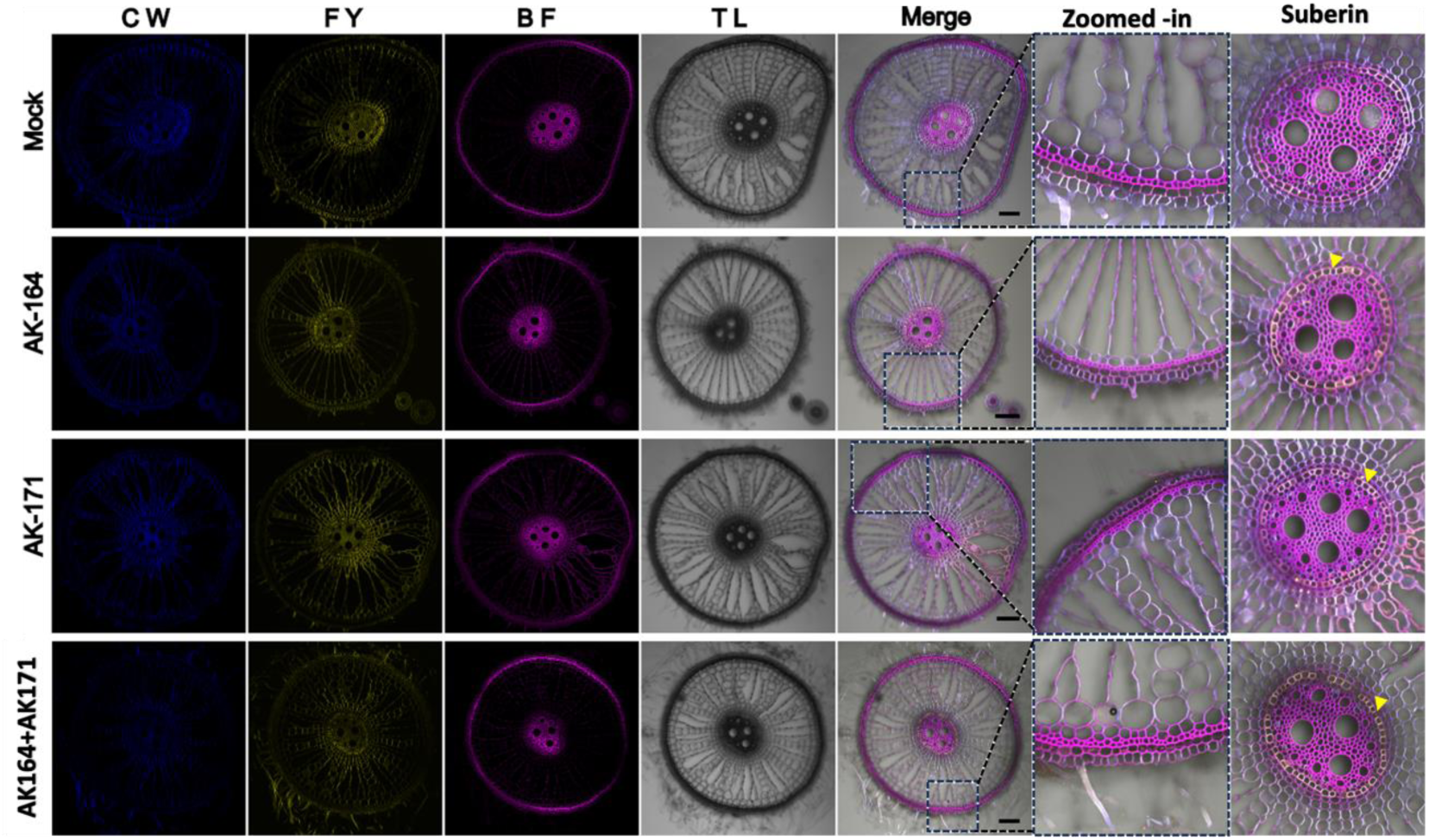
AK164, AK171, and SynCom enhance lignin, suberin, and sclerenchyma formation in rice roots. Root sections of plants colonized with MOCK, AK164, AK171, or SynCom (AK164+AK171) were grown under hydroponic conditions. Multi-serrate sclerenchyma (Zoomed images) in AK164+AK171 roots and enhanced suberin deposition in the endodermal layer (last panel, yellow arrowheads indicate suberin deposition). Panels (left to right) show Calcofluor white (CW), Fluorol Yellow (FY), and Basic Fuchsin (BF), transmission light (TL), and merged channel images. Changes in the sclerenchyma structure are shown in the zoomed-in panel. Bar = 100 mM.

In rice plants grown in hydroponic saline conditions, we observed the collapse of root structure in non-colonized plants that was absent in AK164, AK171, or SynCom plants (Figure 8). Since enhanced sclerenchyma, lignin, and suberin deposition can strengthen organ structure, enhanced robustness of the roots of colonized rice plants may be a reasonable explanation for this finding.

**Figure 8.**
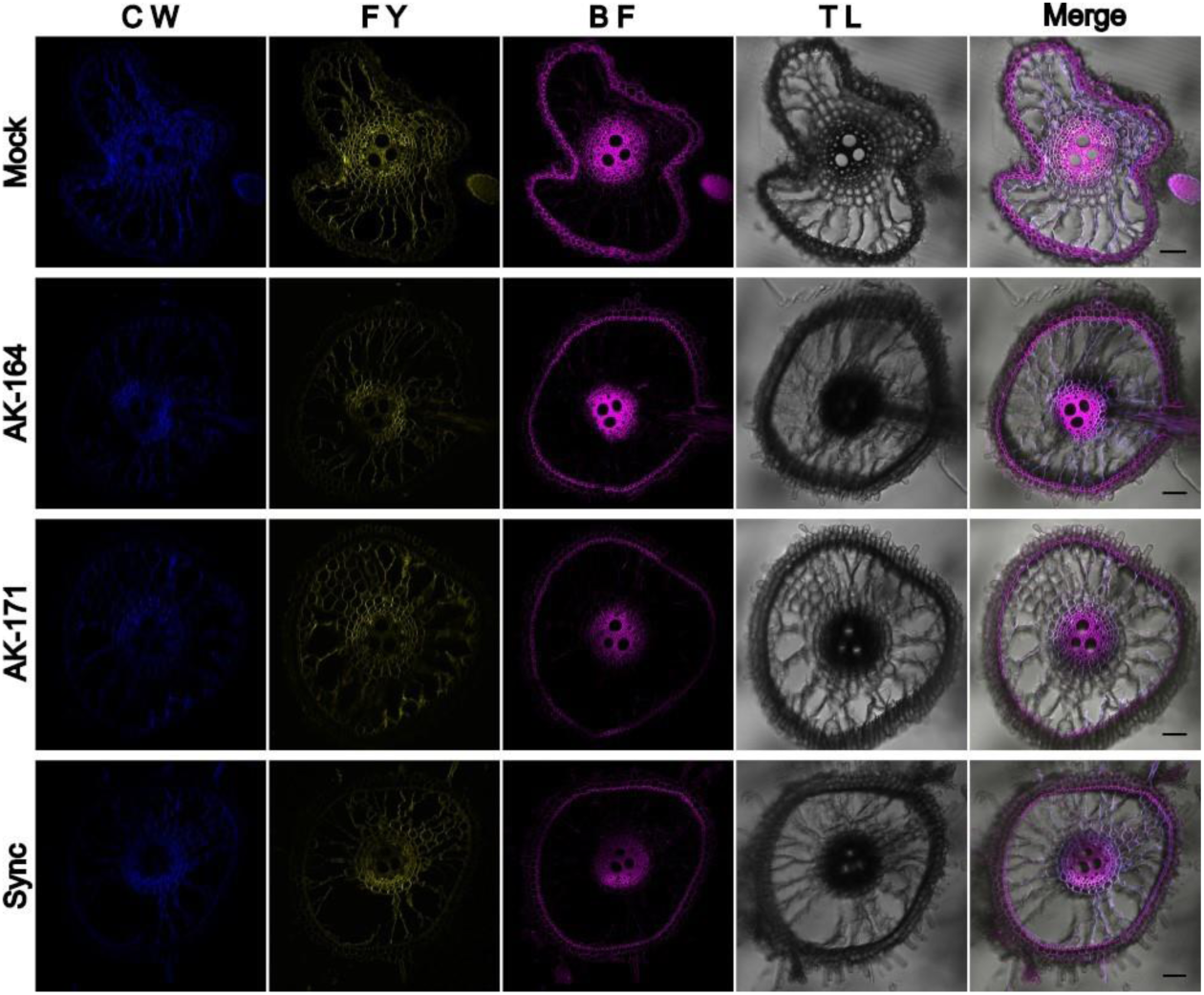
Collapse of rice roots under hydroponic saline conditions is avoided in AK164, AK171, and SynCom colonized plants. Root sections of *O. sativa* cv. Nipponbare colonized with Mock shows structural collapse that is not seen in AK164, AK171, or SynCom colonized plants. Triple staining was performed with Calcofluor white (CW), Fluorol Yellow (FY), and Basic Fuchsin (BF). Transmission light (TL) and merged channel images are shown as indicated.

## Discussion

Given the massive global challenges for food security in the coming decades, securing yield is of primary importance in agriculture. Among these challenges, weather pattern changes result in drought, heat waves, and flooding. Whereas flooding of plants induces hypoxia that can result in damage and death in some species, other plant species can resist hypoxia and survive or even strive under these conditions. The genetic basis of this plasticity has been well studied, and several hormonal pathways and genes have been identified that convey flooding resilience [6, 21–24]. However, little is known whether plant-associated microbes can enhance flooding resilience in sensitive plant species. We show that several bacterial endophytes derived from constantly flooding-challenged mangroves can enhance flooding tolerance in Arabidopsis and rice. Since mangroves thrive in seawater, we also tested whether these microbes could enhance glycophytes salt tolerance. The strains that enhanced flooding tolerance also enhanced salt tolerance in Arabidopsis and rice.

Sixteen strains were analyzed for their capacity to enhance Arabidopsis plant growth on ½ MS plates under standard conditions. To mimic the hypoxic conditions naturally occurring in the natural habitat of the mangrove ecosystem, *A. thaliana* seedlings were then grown using a submerged block method. Among these, AK164 and AK171 were selected as the best performers of rice-inoculated seedlings under normal and saline conditions. *Isoptericola* sp. AK164 enhances the growth of inoculated rice seedlings compared to non-inoculated plants by significantly increasing the shoot DW, root DW, and number of tillers per plant in normal hydroponic conditions. In saline hydroponic conditions, the same parameters were significantly increased compared to mock-inoculated plants. KLBMP 4942 (JX993798) enhanced *Limonium sinense* seed germination by 85 % under normal and 43% under saline conditions while significantly increasing the leaf area under saline conditions, potentially through ACC deaminase production and flavonoid accumulation by the beneficial microbes [25].

To test the ability of AK164 and AK171 to improve plant performance synergistically, we generated a SynCom inoculum of the AK164 and AK171 bacterial strains. Compared to non-inoculated rice plants, we found that SynCom enhanced waterlogging tolerance, yield performance, biomass, and the number of tillers. Under hydroponic saline conditions, SynCom enhanced salt tolerance and yield in shoot DW, root DW, and the number of tillers, which revealed an additive beneficial influence on plant growth under normal hydroponic and hydroponic saline conditions. The additional beneficial effects might be due to complementing each other’s ability to form biofilms, produce secondary metabolites, or stabilize colonization of the host plants [26].

In addition, compared to non-colonized rice plants, agronomic and yield parameters significantly increased with either of the strains but to higher levels for the SynCom under both normal and saline soil in greenhouse conditions. Importantly, the life cycle of rice plants was found to be significantly shortened by colonization of either of the two strains, particularly by the SynCom. Our work confirms other studies showing that applying a SynCom enhances plant growth and yield under greenhouse conditions and in the field [27, 28].

Although it is very important to identify beneficial microbial strains, it is even more important to understand their modus operandi. We tried to find a conserved pattern in analyzing the rice gene expression patterns in response to two taxonomically different mangrove microbes. Since mangrove strains induced rice tolerance to flooding and salt stress as well as enhanced early flowering and yield, we reasoned that the two strains might act by targeting the same signaling pathways in rice plants. The comparative transcriptome analysis of colonized to non-colonized rice roots under hydroponic and hydroponic saline conditions suggested an important role of several genes (JMJ30, NRT1/AIT, SAPK9, and CCD1). JMJ30, a key epigenetic regulator of histone H3 K27 methylation, was identified as the most highly upregulated common gene. The early flowering onset of the inoculated plants under both conditions of our study might be explained by the upregulation of Jumonji JMJ30 [30]. In Arabidopsis, JMJ30 is the key determinant of circadian rhythm. It regulates the flowering time cooperatively with a central oscillator, circadian clock associated1 [31], and late elongated hypocotyl [31, 32], thereby modulating the expression of certain regulators involved in abiotic stress [33]. In Arabidopsis, the circadian clock regulates various processes such as gene expression, photoperiodic flowering, and leaf movement[34]. Along with other factors, JMJ30 might be a determinant of early flowering in OsMATE transgenic plants [33]. JMJ30 promotes callus proliferation by binding to and activating a subset of *LATERAL ORGAN BOUNDARIES DOMAIN* (*LBD*) gene and primarily demethylate trimethylation at lysine 9 of histone H3 (H3K9me3) at *LBD16* and *LBD29*, rather than H3K27me3 [35]. Nitrogen (N) is the most limiting nutrient for crops. Waterlogging causes losses in soil N through denitrification and nitrate leaching, thereby reducing the N pool available for plants and compromising plant growth. In the same context, the nitrogen transporter gene (NRT1) was upregulated in the roots of AK164-and AK171-inoculated plants under hydroponic normal and saline conditions, resulting in a significant increase of plant biomass as well as earlier flowering onset of the inoculated plants. NRT1 is activated by the flowering transcription factor N-mediated heading date-1 (Nhd1), which regulates flowering time. It was also found that Nhd regulates nitrogen uptake and the root biomass in the field [36]. The Stress-Activated Protein Kinase SAPK9, was also upregulated in inoculated plants under hydroponic normal and saline conditions. SAPK9 is a member of the SnRK2 family of protein kinases and one of the core components of the ABA signaling pathway. It facilitates ABA-regulated flowering in rice [37]. SAPK9 also improves drought tolerance and grain yield in rice by modulating cellular osmotic potential, stomatal closure, and stress-responsive gene expression [38]. SAPK9 functions by phosphorylating ABA-responsive element binding protein (AREB) transcriptional factors, including bZIPs (basic region-leucine zipper), to participate in several biological processes, including rice flowering resulting from SAPK/bZIP interactions. Interestingly, SAPK9 binds to bZIP77/OsFD1, a key rice flowering regulator [37], and also interacts with OsMADS23, a positive regulator of osmotic stress, to transcriptionally activate OsNCED2, OsNCED3, OsNCED4, and OsP5CR, which are key components for ABA and proline biosynthesis, respectively. SAPK9 phosphorylation increases the stability and transcriptional activity of OsMADS23, thereby regulating ABA biosynthesis and providing a novel strategy to improve salinity and drought tolerance in rice[39]. In rice, ABA regulates suberin deposition via *TREHALOSE-PHOSPHATE-SYNTHASE* (OsTPS8)[40]. Over-expression of *OsTPS8* enhances salinity tolerance without yield penalty. In contrast, the *ostps8* mutant showed significantly reduced soluble sugars, Casparian bands, and suberin deposition in the roots compared to the WT and overexpression lines [40]. This might explain the suberin deposition in the cell walls of the root cells in the PGPB-inoculated plants in our study (Figure 5-13). ABA can also regulate the lignin biosynthesis by phosphorylating the master lignin transcription factor NST1 – a substrate for ABA-dependent SnRK2 kinases, suggesting a tight control of ABA signaling in secondary cell wall deposition [41].

Under hydroponic conditions, the roots of inoculated plants upregulated carotenoid cleavage dioxygenases (CCD1). CCDs are important biosynthetic enzymes for distinct carotenoids. A ccd1 ccd4 mutant showed fewer meristematic cell divisions and less lateral root branching compared to mock roots [42], which can be rescued by root indigenous β-cyclocitral. In addition, *OsCCD4a*-RNAi-*7* lines showed increases in the leaf carotenoids α-carotene, β-carotene, lutein, antheraxanthin, and violaxanthin relative to non-transgenic plants. Interestingly, seeds of *OsCCD1*-RNAi-*8* plants displayed a 1.4-fold increase in total carotenoids [43]. In contrast, inoculated plant roots exhibited upregulation of the CAROTENOID CLEAVAGE DIOXYGENASE 8 (CCD8) gene. This gene plays a role in the biosynthesis of the phytohormone strigolactone, which enhances tomato yield production [44]. *OsCCD8b* regulates rice (*Oryza sativa*) tillering, whereas *ZmCCD8* plays essential roles in root and shoot development, with smaller roots, shorter internodes, and longer tassels in *Zmccd8* mutant plants [45, 46].

Abiotic stress, including flooding and salinity, drastically inhibits plant growth, and productivity is severely reduced [47, 48]. Multiple mechanisms are used to restore cellular homeostasis and promote survival [49]. Osmotic stresses, including salt, drought, and/or PEG treatment, have been shown to affect plant cell water balance. For example, the root water uptake capacity (i.e., root hydraulic conductivity) was inhibited by salinity [50, 51]. Transcriptome analysis of the roots of rice plants under both hydroponic and saline conditions showed several aquaporins (AQPs) to be downregulated upon bacterial inoculation using AK164 and AK171. The aquaporins are membrane proteins of plasma membranes (PIPs) or the tonoplast (TIPs). Aquaporins are also thought to be involved in stress-coping mechanisms, such as altering the hydraulic conductivity of tissues [52, 53]. In rice (*Oryza sativa* L. cv. Nipponbare), 33 aquaporins were identified, including 11 PIPs, 10 TIPs, 10 NIPs and 2 SIPs [54]. Healthy plants must have a constant water supply, which is important for growing tissues to maintain turgor pressure upon cell enlargement [55]. AK164 and AK171 inoculated plants showed downregulation of a few genes, of which the AQPs represented the only family of related functions. The concreted regulation of PIP1;2, PIP2;3, PIP2-4, and PIP2;5, suggest a regulation of the plant water status by the two microbial strains. Various studies suggest that a low abundance of AQP proteins reduces water permeability, and a high abundance (over-expression) increases the hydraulic conductivity of biological membranes [56]. It was found that plants with mid-to low-level overexpression of OsPIP1;1 showed salt resistance, [57]implicating that OsPIP1;1 plays an important role in regulating water homeostasis *in planta*. Although no significant changes in PIP1;1 expression was observed in our study under salinity, we observed significant changes in PIP1;2 under normal conditions of the PGPB-inoculated roots. OsPIP2;1 decreased root length, surface area, root volume, and root tip number [58]. Additionally, PIP1;1 has high water channel activity when co-expressed with PIP2, and that PIP1–PIP2 random hetero tetramerization not only allows PIP1;1 to arrive at the plasma membrane but also results in enhanced activity of PIP2;1 [59]. These results suggest the importance of both PIP1 and PIP2 aquaporins for controlling water movement across the plasma membrane [60]. In agreement with our findings, the overexpression of *OsPIP1;1* at very high levels markedly decreases fertility, as we found in MOCK plants in normal conditions, whereas gene expression at low or medium levels raises seed yield but does not affect single-grain weight [57]. This was implemented by suggesting the role of OsPIP1;1 as a seed set, probably by affecting pollen germination in the stigma or pollen tube growth [61]. Several authors reported that *PIP* expression responded to salt and drought stress in *Arabidopsis* and rice [62–64]. The expression pattern of a particular aquaporin gene may vary within or across plant species. This variability can be rooted in the different cultivars, developmental stages, or stress conditions. In all, it has been stated that TIPs and PIPs together may help the plant adjust water transport through transcriptional regulations to tune the water balance under stress [65, 66]. In *Arabidopsis* and rice, the expression response to dehydration and salinity in some *PIP*s was mediated by ABA [64, 67, 68]. These data indicated that, as a stress-signaling molecule, ABA is involved in regulating some *TIP*s and *PIP*s expression in response to abiotic stresses. It was also reported that the expression of rice *TIP*s was significantly upregulated by dehydration, salinity, and ABA treatments. It was found that TIP1 expression in shoots and roots of rice seedlings was influenced by water stress, salt stress, and exogenous ABA [69].

In addition to water transport, AQPs may contribute to hypoxia tolerance by transporting O_2_ [55], H_2_O_2_, and lactic acid. The complex responses toward oxygen deprivation may involve ethylene, abscisic acid, or other hormonal factors and signaling molecules [70].

## Conclusion

At present, it becomes clear that PGPB strains have highly conserved mechanisms for interacting with plants to enhance stress resilience. The underlying mechanisms slowly emerge and suggest that many beneficial mechanisms are based on reprogramming the plant genomes. Importantly, the beneficial microbes not only enhance the stress tolerance but also the growth and yield of various crops. Our data unanimously show that two mangrove strains, *Isoptericola* AK164, and *Tritonibacter mobilis* AK171, along with their SynCom, can enhance the resilience of monocot and dicot plants to waterlogging and salinity stress without any growth and yield penalty. Overall, our results highlight the potential of symbiotic bacteria as an eco-friendly innovation towards sustainable agriculture in the face of the challenges of climate change.

## Supplementary Figures

**Figure S1:**
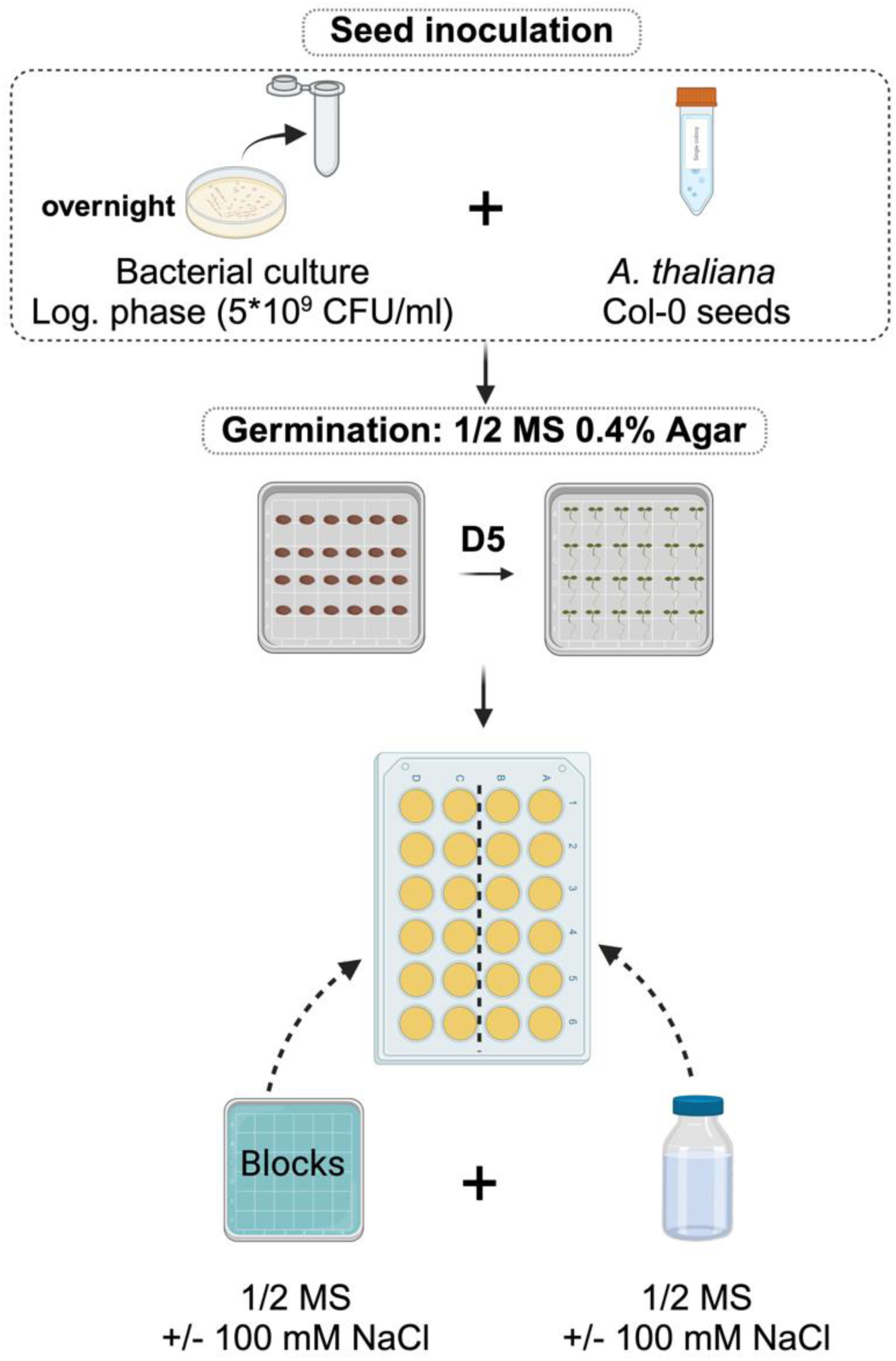
Screen for potential PGPBs of the isolated mangrove bacterial collection in *A. thaliana*.

**Figure S2:**
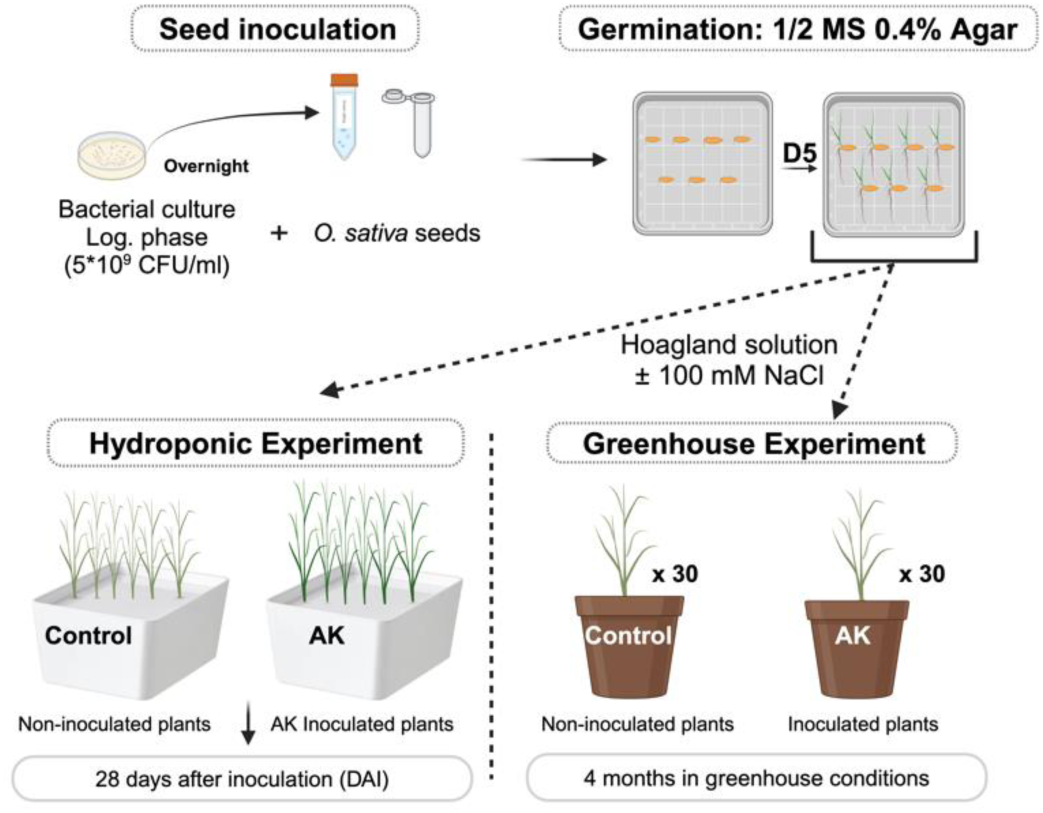
Workflow of the hydroponic and greenhouse experiments using modified Hoagland nutrient solution.

**Figure S3:**
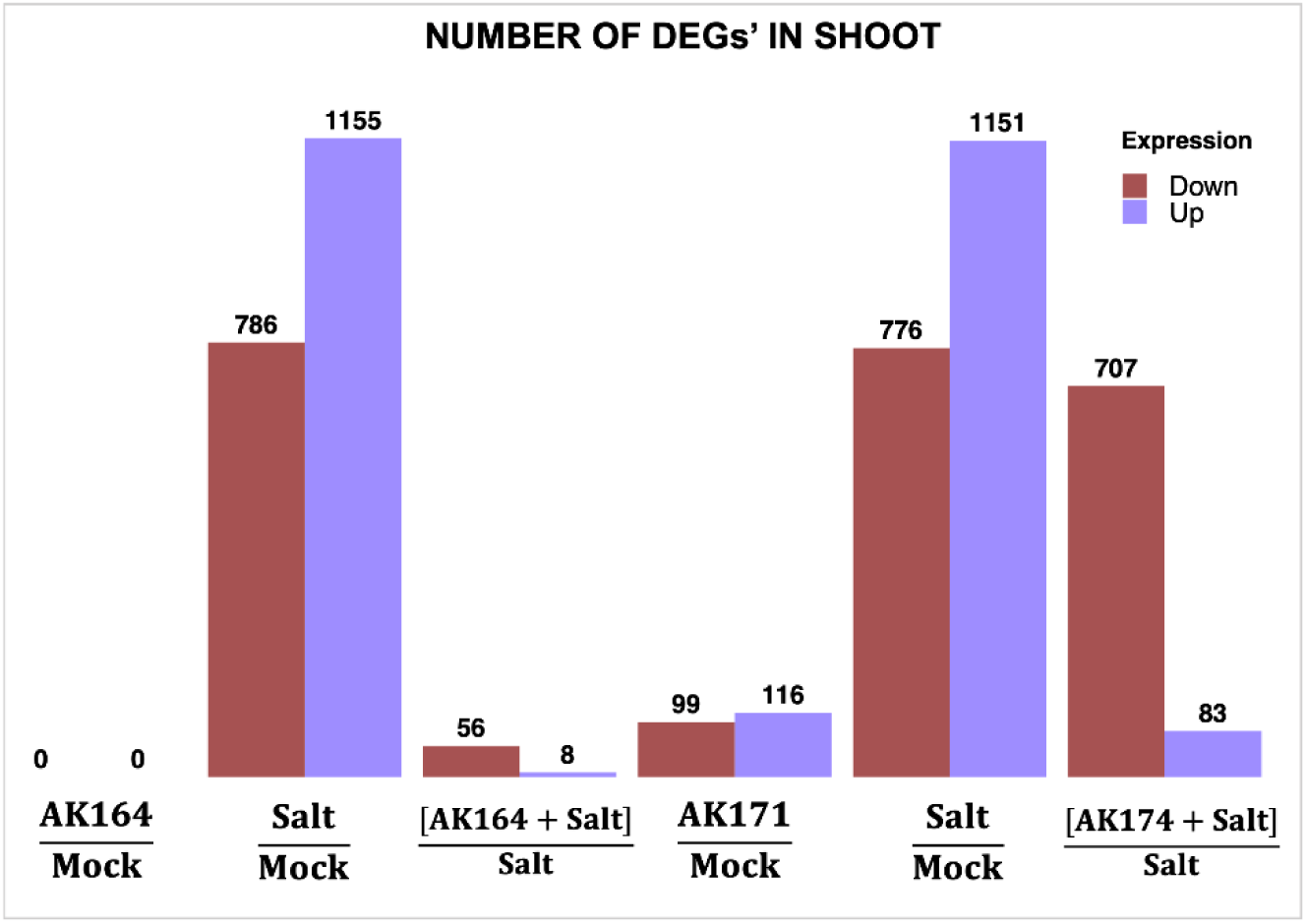
Shoot RNA sequencing data. DEGs of shoots of rice plants grown under normal and saline hydroponic conditions stress treated with AK164, AK171, or MOCK inoculated. Number of DEGs (log 2 > 1, P < 0.01) in rice shoots colonized with AK164, AK171, or MOCK under normal (0 mM NaCl) and saline (100 mM NaCl) hydroponic conditions.

**Supplementary Table 1:** Beneficial impact of the best-performing mangrove isolates on the shoot and root systems of *O. sativa* seedlings (28 days old) under normal hydroponic conditions (Hoagland nutrient solution). AK031, B. seohaeanensis; AK073, Thalassospira tepidiphila; AK116, H. locisalis; AK157, Demequina activiva, AK164, I. chiayiensis; AK171, T. mobilis; AK181, P. azotoformans; AK255, I. chiayiensis; and AK266, Paenibacillus lautus. Different letters indicate statistically significant differences depending on Dunn’s multiple comparison test (P < 0.05). Identical letters indicate no significant differences at P 0.05.

| NO. | Strain | SHOOT FRESH WEIGHT (GM%) |  | LENGTH (CM%) |  | REPRODUCTION |
| --- | --- | --- | --- | --- | --- | --- |
|  |  | Shoot | Root | Shoot | Root | NO. OF TILLERS |
| 1 | AK031 | 8.98<sup>ab</sup> | 5.18<sup>bc</sup> | 22.88<sup>a</sup> | 9.60<sup>b</sup> | 48.13<sup>BC</sup> |
| 2 | AK073 | 25.48<sup>ab</sup> | 70.76<sup>a</sup> | 20.71<sup>a</sup> | 47.40<sup>a</sup> | 40.76<sup>CD</sup> |
| 3 | AK116 | 28.85<sup>ab</sup> | 71.42<sup>a</sup> | 4.88<sup>b</sup> | 6.19<sup>b</sup> | 40.76<sup>CD</sup> |
| 4 | AK157 | 31.04<sup>ab</sup> | 71.93<sup>a</sup> | 4.66<sup>b</sup> | 7.04<sup>b</sup> | 100.00<sup>A</sup> |
| 5 | AK164 | 37.59<sup>a</sup> | 80.38<sup>a</sup> | 3.19<sup>b</sup> | 15.81<sup>ab</sup> | 96.31<sup>A</sup> |
| 6 | AK171 | 40.49<sup>a</sup> | 73.65<sup>a</sup> | 4.88<sup>b</sup> | 14.17<sup>b</sup> | 88.89<sup>AB</sup> |
| 7 | AK181 | 44.32<sup>a</sup> | 80.55<sup>a</sup> | 11.59<sup>a</sup><br>b | 26.05<sup>ab</sup> | 77.78<sup>ABC</sup> |
| 8 | AK255 | 31.14<sup>A</sup><br>B | 51.61<sup>A</sup><br>B | 21.68<br>A | 27.90<sup>AB</sup> | 74.09<sup>ABC</sup> |

**Supplementary Table 2:** Beneficial impact of the best-performing mangrove isolates on the shoot and root systems of O. sativa seedlings (28 days old) under saline hydroponic conditions (Hoagland solution with 100mM NaCl). AK031, B. seohaeanensis; AK073, T. tepidiphila; AK116, H. locisalis; AK157, D. activiva, AK164, I. chiayiensis; AK171, T. mobilis; AK181, P. azotoformans; AK255, I. chiayiensis; and AK266, Paenibacillus lautus. Different letters indicate statistically significant differences depending on Dunn’s multiple comparison test (P < 0.05). Identical letters indicate no significant differences at P 0.05.

| No. | Strain | Shoot Fresh Weight (mg%) |  | Length (cm%) |  | Reproduction |
| --- | --- | --- | --- | --- | --- | --- |
|  |  | Shoot | Root | Shoot | Root | No. of Tillers |
| 1 | AK031 | 60.73<sup>bc</sup> | 103.01<sup>cd</sup> | 0.49<sup>ab</sup> | -3.81<sup>e</sup> | 3.73<sup>b</sup> |
| 2 | AK073 | 53.24<sup>bcd</sup> | 53.25<sup>de</sup> | 28.46<sup>ab</sup> | 10.08<sup>bc</sup> | -3.73<sup>b</sup> |
| 3 | AK116 | 52.42<sup>bcd</sup> | 41.88<sup>de</sup> | 29.61<sup>ab</sup> | 21.59<sup>a</sup> | -3.73<sup>b</sup> |
| 4 | AK157 | 24.57<sup>cd</sup> | 11.91<sup>e</sup> | 10.85<sup>c</sup> | -2.70<sup>de</sup> | 0.00<sup>b</sup> |
| 5 | AK164 | 87.12<sup>ab</sup> | 284.29<sup>a</sup> | 38.91<sup>a</sup> | 20.32<sup>ab</sup> | 48.18<sup>a</sup> |
| 6 | AK171 | 116.80<sup>a</sup> | 193.58<sup>b</sup> | 37.18<sup>ab</sup> | 21.35<sup>a</sup> | 33.33<sup>ab</sup> |
| 7 | AK181 | 33.61<sup>cd</sup> | 45.33<sup>de</sup> | 12.70<sup>c</sup> | 6.98<sup>cd</sup> | 7.38<sup>b</sup> |
| 8 | AK255 | 39.27<sup>cd</sup> | 144.71<sup>bc</sup> | 13.86<sup>c</sup> | -18.57<sup>f</sup> | 14.84<sup>ab</sup> |

**Supplementary Table 3:**
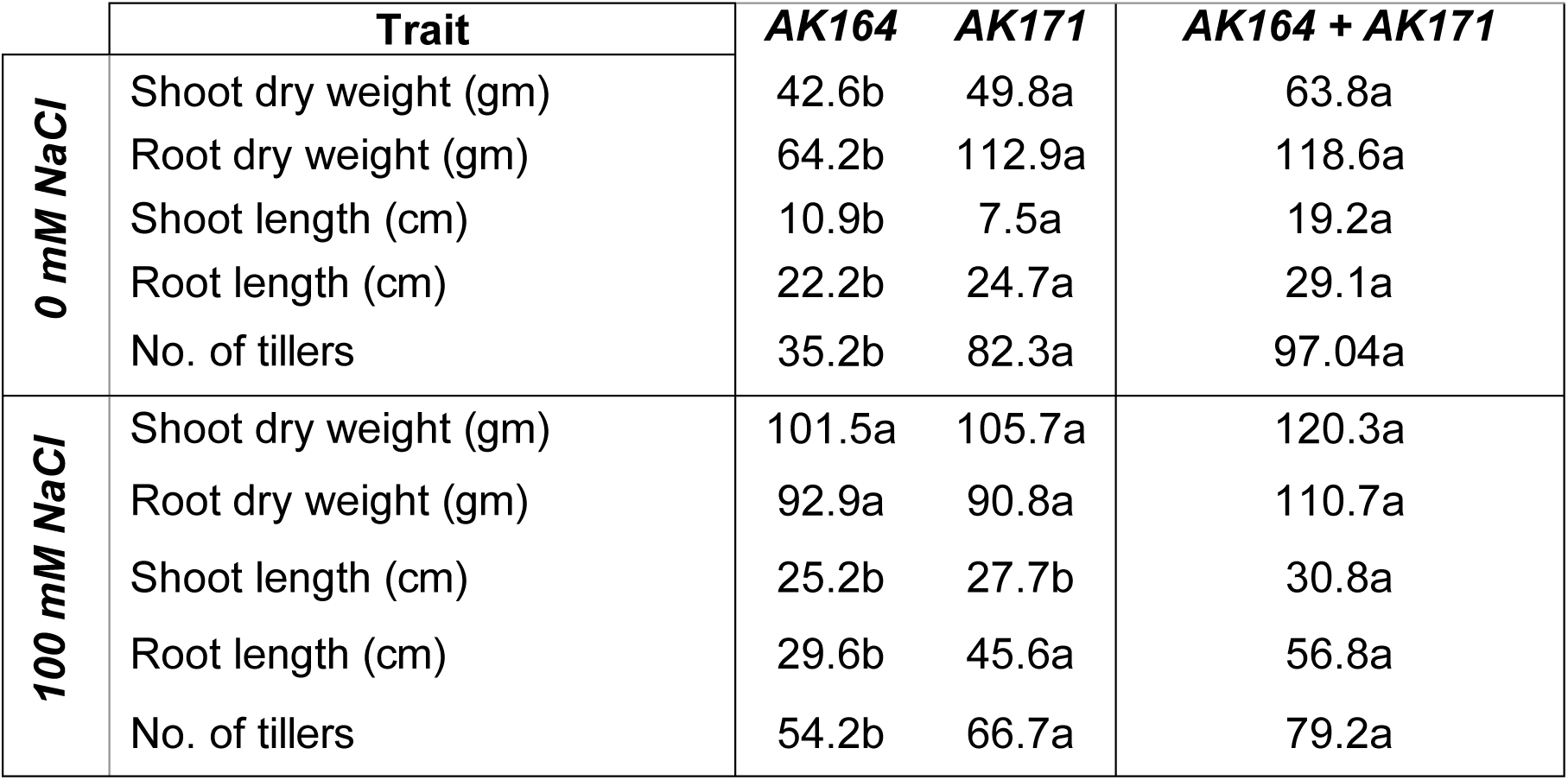
Beneficial increase (%) of various traits in AK164, AK171, and SynCom (AK164+AK171) colonized *O. sativa* in hydroponic growth (0 or 100 mM NaCl) compared to non-inoculated plants. Different letters indicate statistically significant differences depending on Dunn’s multiple comparison test (P < 0.05). Identical letters indicate no significant differences at *p = 0.05*.

**Supplementary Table 4:**
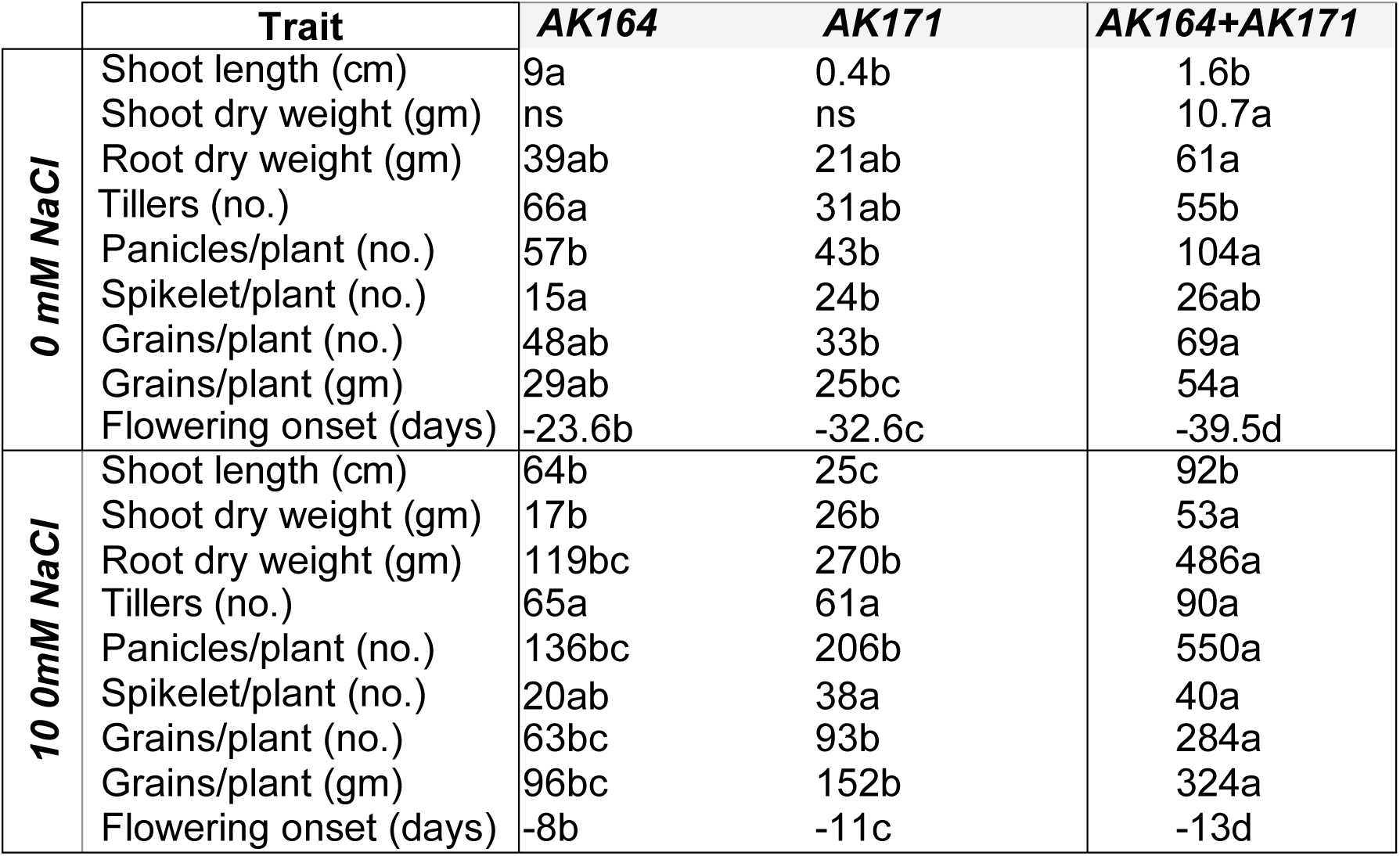
Beneficial increase (%) of various traits in AK164, AK171, and SynCom (AK164+AK171) colonized *O. sativa* growing in soil under greenhouse conditions compared to non-inoculated plants. Different letters indicate statistically significant differences depending on Dunn’s multiple comparison test (P < 0.05). Identical letters indicate no significant differences at P 0.05.

**Supplementary Table 5:**
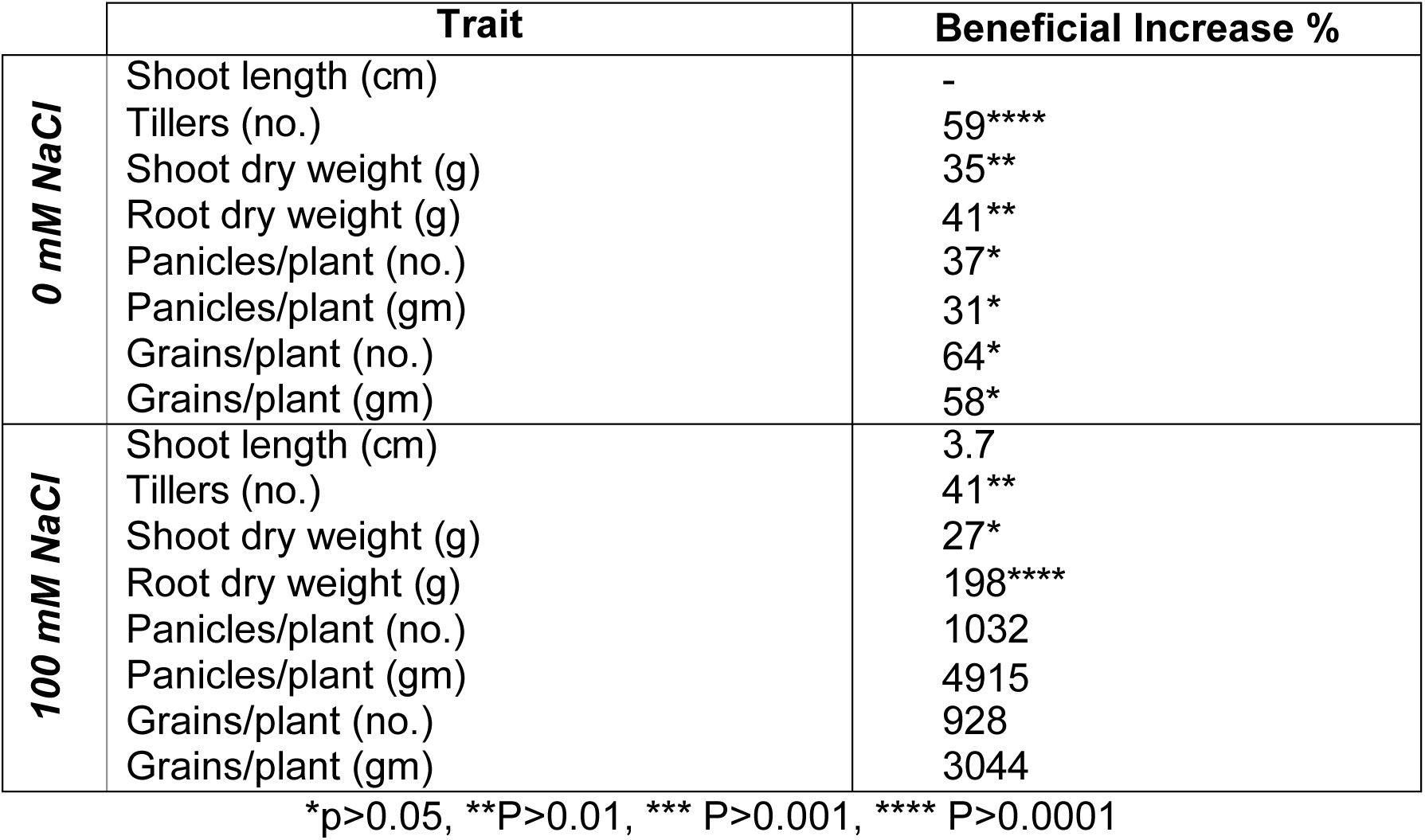
Beneficial increase (%) of various traits and yield in SynCom (AK164+AK171) colonized *O. sativa* plants growing on soil (0 or 100 mM NaCl) under greenhouse conditions compared to non-inoculated plants.

|  | Trait | Beneficial Increase % |
| --- | --- | --- |
| **0 mM NaCl** | Shoot length (cm) | - |
|  | Tillers (no.) | 59**** |
|  | Shoot dry weight (g) | 35** |
|  | Root dry weight (g) | 41** |
|  | Panicles/plant (no.) | 37* |
|  | Panicles/plant (gm) | 31* |
|  | Grains/plant (no.) | 64* |
|  | Grains/plant (gm) | 58* |
| **100 mM NaCl** | Shoot length (cm) | 3.7 |
|  | Tillers (no.) | 41** |
|  | Shoot dry weight (g) | 27* |
|  | Root dry weight (g) | 198**** |
|  | Panicles/plant (no.) | 1032 |
|  | Panicles/plant (gm) | 4915 |
|  | Grains/plant (no.) | 928 |
|  | Grains/plant (gm) | 3044 |

\*p>0.05, \*\*P>0.01, \*\*\* P>0.001, \*\*\*\* P>0.0001

## Notes

https://www.ncbi.nlm.nih.gov/geo/query/acc.cgi?acc=GSE269165

### Competing Interest Statement

The authors have declared no competing interest.

### Summary of Updates

Author names were corrected as they were not in the correct order and style

## References

1. FAO, Leveraging food systems for inclusive rural transformation., in The State of Food and Agriculture (SOFA) 2017. 2017: Rome. p. #181.

2. FAO, FAOSTAT. Online statistical database. 2017.

3. Lee, S.C., et al., Molecular characterization of the submergence response of the Arabidopsis thaliana ecotype Columbia. New Phytol, 2011. 190(2): p. 457–71.

4. Hsu, F.C., et al., Insights into hypoxic systemic responses based on analyses of transcriptional regulation in Arabidopsis. PLoS One, 2011. 6(12): p. e28888.

5. Jackson, M.B. and P.C. Ram, Physiological and molecular basis of susceptibility and tolerance of rice plants to complete submergence. Ann Bot, 2003. 91 **Spec No**(2): p. 227–41.

6. Yeung, E., et al., A stress recovery signaling network for enhanced flooding tolerance in Arabidopsis thaliana. Proc Natl Acad Sci U S A, 2018. 115(26): p. E6085–E6094.

7. Tsai, K.J., et al., Ethylene-Regulated Glutamate Dehydrogenase Fine-Tunes Metabolism during Anoxia-Reoxygenation. Plant Physiol, 2016. 172(3): p. 1548–1562.

8. Fukao, T., E. Yeung, and J. Bailey-Serres, The submergence tolerance regulator SUB1A mediates crosstalk between submergence and drought tolerance in rice. Plant Cell, 2011. 23(1): p. 412–27.

9. Mustroph, A., et al., Cross-kingdom comparison of transcriptomic adjustments to low-oxygen stress highlights conserved and plant-specific responses. Plant Physiol, 2010. 152(3): p. 1484–500.

10. Juntawong, P., et al., Ribosome profiling: a tool for quantitative evaluation of dynamics in mRNA translation.Methods. 2015. 1284: p. 139–173.

11. Tsuji, H., et al., Dynamic and reversible changes in histone H3-Lys4 methylation and H3 acetylation occurring at submergence-inducible genes in rice. Plant Cell Physiol, 2006. 47(7): p. 995–1003.

12. Van Zelm, E., Y. Zhang, and C. Testerink, Salt Tolerance Mechanisms of Plants. Annual Review of Plant Biology, 2020. 71: p. 403–433.

13. Romano, S., F. Vanni, and D. Viaggi, Economics of culture and food in evolving agri-food systems and rural areas. Bio-based and Applied Economics, 2020. 9(2): p. 127–136.

14. Saad, M.M., A.A. Eida, and H. Hirt, Tailoring plant-associated microbial inoculants in agriculture: a roadmap for successful application. J Exp Bot, 2020. 71(13): p. 3878–3901.

15. Alghamdi, A.K., et al., Complete genome sequence analysis of plant growth-promoting bacterium, Isoptericola sp. AK164 isolated from the rhizosphere of Avicennia marina growing at the Red Sea coast. Arch Microbiol, 2023. 205(9): p. 307.

16. Kawahara Y, et al., Improvement of the Oryza sativa Nipponbare reference genome using next generation sequence and optical map data. Rice, 2013. 6(1): p. 4.

17. Sakai, H., et al., Rice Annotation Project Database (RAP-DB): an integrative and interactive database for rice genomics. Plant Cell Physiol, 2013. 54(2): p. e6.

18. Pertea, M., et al., StringTie enables improved reconstruction of a transcriptome from RNA-seq reads. Nat Biotechnol, 2015. 33(3): p. 290–5.

19. Sexauer, M., et al., Visualizing polymeric components that define distinct root barriers across plant lineages. Development, 2021. 148(23).

20. Kruskal, W.H. and W.A. Wallis, Use of Ranks in One-Criterion Variance Analysis. Journal of the American Statistical Association, 1952. 47(260): p. 583–621.

21. Jhan, L.H., et al., Integrative pathway and network analysis provide insights on flooding-tolerance genes in soybean. Scientific Reports, 2023. 13(1): p. 1–22.

22. Singh, T., et al., The hidden harmony: Exploring ROS-phytohormone nexus for shaping plant root architecture in response to environmental cues. Plant Physiol Biochem, 2024. 206: p. 108273.

23. Khan, N., et al., Crosstalk amongst phytohormones from planta and PGPR under biotic and abiotic stresses. Plant Growth Regulation, 2020. 90(2): p. 189–203.

24. Sasidharan, R., et al., Signal dynamics and interactions during flooding stress. Plant Physiology, 2018. 176(2): p. 1106–1117.

25. Qin, S., et al., Isolation of ACC deaminase-producing habitat-adapted symbiotic bacteria associated with halophyte Limonium sinense (Girard) Kuntze and evaluating their plant growth-promoting activity under salt stress. Plant and Soil, 2014. 374(1): p. 753–766.

26. Shayanthan, A., P.A.C. Ordoñez, and I.J. Oresnik, The Role of Synthetic Microbial Communities (SynCom) in Sustainable Agriculture. Frontiers in Agronomy, 2022. 4.

27. Nadeem, S.M., et al., The role of mycorrhizae and plant growth promoting rhizobacteria (PGPR) in improving crop productivity under stressful environments. Biotechnol Adv, 2014. 32(2): p. 429–48.

28. Duarte, B., et al., Marine Plant Growth Promoting Bacteria (PGPB) inoculation technology: Testing the effectiveness of different application methods to improve tomato plants tolerance against acute heat wave stress. Plant Stress, 2024. 11.

29. Xu, K., et al., Sub1A is an ethylene-response-factor-like gene that confers submergence tolerance to rice. Nature, 2006. 442(7103): p. 705–8.

30. Kuroda, T. and T. Tsuchiya, Multidrug efflux transporters in the MATE family. Biochim Biophys Acta, 2009. 1794(5): p. 763–8.

31. Lai, A.G., et al., CIRCADIAN CLOCK-ASSOCIATED 1 regulates ROS homeostasis and oxidative stress responses. Proc Natl Acad Sci U S A, 2012. 109(42): p. 17129–34.

32. Lu, S.X., et al., The Jumonji C domain-containing protein JMJ30 regulates period length in the Arabidopsis circadian clock. Plant Physiol, 2011. 155(2): p. 906–15.

33. Tiwari, M., et al., Expression of OsMATE1 and OsMATE2 alters development, stress responses and pathogen susceptibility in Arabidopsis. Sci Rep, 2014. 4: p. 3964.

34. McClung, C.R., McClung, C. R. Plant circadian rhythms. Plant Cell, 2006(18): p. 792–803.

35. Lee, K., O.S. Park, and P.J. Seo, JMJ30-mediated demethylation of H3K9me3 drives tissue identity changes to promote callus formation in Arabidopsis. Plant J, 2018. 95(6): p. 961–975.

36. Li, K., et al., The rice transcription factor Nhd1 regulates root growth and nitrogen uptake by activating nitrogen transporters. Plant Physiol, 2022. 189(3): p. 1608–1624.

37. Liu, X., et al., Protein Interactomic Analysis of SAPKs and ABA-Inducible bZIPs Revealed Key Roles of SAPK10 in Rice Flowering. Int J Mol Sci, 2019. 20(6).

38. Dey, A., et al., The sucrose non-fermenting 1-related kinase 2 gene SAPK9 improves drought tolerance and grain yield in rice by modulating cellular osmotic potential, stomatal closure and stress-responsive gene expression. BMC Plant Biol, 2016. 16(1): p. 158.

39. Li, X., et al., OsMADS23 phosphorylated by SAPK9 confers drought and salt tolerance by regulating ABA biosynthesis in rice. PLoS Genet, 2021. 17(8): p. e1009699.

40. Vishal, B., et al., OsTPS8 controls yield-related traits and confers salt stress tolerance in rice by enhancing suberin deposition. New Phytol, 2019. 221(3): p. 1369–1386.

41. Liu, C., et al., Abscisic acid regulates secondary cell-wall formation and lignin deposition in Arabidopsis thaliana through phosphorylation of NST1. Proceedings of the National Academy of Sciences, 2021. 118(5).

42. Dickinson, A.J., et al., *beta-Cyclocitral is a conserved root growth regulator*. Proc Natl Acad Sci U S A, 2019. 116(21): p. 10563–10567.

43. Ko, M.R., et al., RNAi-mediated suppression of three carotenoid-cleavage dioxygenase genes, OsCCD1, 4a, and 4b, increases carotenoid content in rice. J Exp Bot, 2018. 69(21): p. 5105–5116.

44. Kohlen, W., et al., The tomato CAROTENOID CLEAVAGE DIOXYGENASE8 (SlCCD8) regulates rhizosphere signaling, plant architecture and affects reproductive development through strigolactone biosynthesis. New Phytol, 2012. 196(2): p. 535–547.

45. Guan, J.C., et al., Diverse roles of strigolactone signaling in maize architecture and the uncoupling of a branching-specific subnetwork. Plant Physiol, 2012. 160(3): p. 1303–17.

46. Arite, T., et al., DWARF10, an RMS1/MAX4/DAD1 ortholog, controls lateral bud outgrowth in rice. Plant J, 2007. 51(6): p. 1019–29.

47. Atkinson, N.J. and P.E. Urwin, The interaction of plant biotic and abiotic stresses: from genes to the field. Journal of Experimental Botany, 2012. 63(10): p. 3523–3543.

48. Osakabe, Y., et al., Response of plants to water stress. Front Plant Sci, 2014. 5: p. 86.

49. Golldack, D., et al., Tolerance to drought and salt stress in plants: Unraveling the signaling networks. Front Plant Sci, 2014. 5: p. 151.

50. Boursiac, Y., et al., Early effects of salinity on water transport in Arabidopsis roots. Molecular and cellular features of aquaporin expression. Plant Physiol, 2005. 139(2): p. 790–805.

51. Martinez-Ballesta, M.C., et al., Different blocking effects of HgCl2 and NaCl on aquaporins of pepper plants. J Plant Physiol, 2003. 160(12): p. 1487–92.

52. Merlaen, B., et al., Physiological responses and aquaporin expression upon drought and osmotic stress in a conservative vs prodigal Fragaria x ananassa cultivar. Plant Physiol Biochem, 2019. 145: p. 95–106.

53. Qiao, Y., et al., Exogenous melatonin alleviates PEG-induced short-term water deficiency in maize by increasing hydraulic conductance. BMC Plant Biol, 2020. 20(1): p. 218.

54. Sakurai, J., et al., Identification of 33 rice aquaporin genes and analysis of their expression and function. Plant Cell Physiol, 2005. 46(9): p. 1568–77.

55. Zwiazek, J.J., et al., Significance of oxygen transport through aquaporins. Sci Rep, 2017. 7: p. 40411.

56. Martre, P., et al., Plasma membrane aquaporins play a significant role during recovery from water deficit. Plant Physiol, 2002. 130(4): p. 2101–10.

57. Liu, C., et al., Aquaporin OsPIP1;1 promotes rice salt resistance and seed germination. Plant Physiol Biochem, 2013. 63: p. 151–8.

58. Ding, L., et al., Aquaporin PIP2;1 affects water transport and root growth in rice (Oryza sativa L.). Plant Physiol Biochem, 2019. 139: p. 152–160.

59. Bienert, M.D., et al., Heterotetramerization of Plant PIP1 and PIP2 Aquaporins Is an Evolutionary Ancient Feature to Guide PIP1 Plasma Membrane Localization and Function. Front Plant Sci, 2018. 9: p. 382.

60. Kudoyarova, G., et al., The Role of Aquaporins in Plant Growth under Conditions of Oxygen Deficiency. Int J Mol Sci, 2022. 23(17).

61. Wang, Y., et al., Versatile Roles of Aquaporins in Plant Growth and Development. Int J Mol Sci, 2020. 21(24).

62. Li, G.W., et al., Transport functions and expression analysis of vacuolar membrane aquaporins in response to various stresses in rice. J Plant Physiol, 2008. 165(18): p. 1879–88.

63. Alexandersson, E., et al., Whole gene family expression and drought stress regulation of aquaporins. Plant Mol Biol, 2005. 59(3): p. 469–84.

64. Guo, L., et al., Expression and functional analysis of the rice plasma-membrane intrinsic protein gene family. Cell Res, 2006. 16(3): p. 277–86.

65. Bailey-Serres, J. and L.A. Voesenek, Flooding stress: acclimations and genetic diversity. Annu Rev Plant Biol, 2008. 59: p. 313–39.

66. Xiong, L. and J. Zhu, Regulation of abscisic acid biosynthesis. Plant Physiol., 2003. 133((1):. PMID: 12970472; PMCID: PMC523868.): p. 29–36.

67. Jang, J.Y., et al., An Expression Analysis of a Gene Family Encoding Plasma Membrane Aquaporins in Response to Abiotic Stresses in Arabidopsis Thaliana. Plant Molecular Biology, 2004. 54: p. 713–725.

68. Lian, H.L., et al., Upland rice and lowland rice exhibited different PIP expression under water deficit and ABA treatment. Cell Res, 2006. 16(7): p. 651–60.

69. Liu, Q., M. Umeda, and H. Uchimiya, Isolation and expression analysis of two rice genes encoding the major intrinsic protein. Plant Molecular Biology, 1994(26): p. 2003–2007.

70. Almahasheer, H., et al., Low Carbon sink capacity of Red Sea mangroves. Scientific Reports, 2017. 7(1): p. 1–10.

